# Reorganization of the neurobiology of language after sentence overlearning

**DOI:** 10.1101/2020.09.11.293167

**Authors:** Jeremy I Skipper, Sarah Aliko, Stephen Brown, Yoon Ju Jo, Serena Lo, Emilia Molimpakis, Daniel R Lametti

## Abstract

There is a widespread assumption that there are a static set of ‘language regions’ in the brain. Yet, people still regularly produce familiar ‘formulaic’ expressions when those regions are severely damaged. This suggests that the neurobiology of language varies with the extent of word sequence learning and might not be fixed. We test the hypothesis that perceiving sentences is mostly supported by sensorimotor regions involved in speech production and not ‘language regions’ after overlearning. Twelve participants underwent two sessions of behavioural testing and functional magnetic resonance imaging (fMRI), separated by 15 days. During this period, they repeated two sentences 30 times each, twice a day. In both fMRI sessions, participants ‘passively’ listened to those two sentences and novel sentences. Lastly, they spoke novel sentences. Behavioural results confirm that participants overlearned sentences. Correspondingly, there was an increase or recruitment of sensorimotor regions involved in sentence production and a reduction in activity or inactivity for overlearned sentences in regions involved in listening to novel sentences. The global network organization of the brain changed by ∼45%, mostly through lost connectivity. Thus, there was a profound reorganization of the neurobiology of speech perception after overlearning towards sensorimotor regions not considered in most contemporary models and away from the ‘language regions’ posited by those models. These same sensorimotor regions are generally preserved in aphasia and Alzheimer’s disease, perhaps explaining residual abilities with formulaic language. These and other results warrant reconsidering static neurobiological models of language.

## Introduction

It is widely assumed that the brain regions supporting speech production, perception and language comprehension are largely spatially fixed. Historical models consider ‘Broca’s area’ the locus of speech production and ‘Wernicke’s area’ the locus of comprehension^1,2^. Popular contemporary models include these left hemisphere inferior frontal gyrus and perisylvian regions and a small number of others, most notably premotor cortex and bilateral superior and middle temporal gyri^3–11^. Illustrating this fixity assumption, various aspects of the inferior frontal gyrus and superior temporal plane are regularly described as ‘the language network’, with the latter phrase appearing in more than 5500 articles on Google Scholar (assessed September, 2020)^8^.

Yet, there is a long history of evidence suggesting that language is more distributed throughout the brain than is belied by these models^12–14^. This is illustrated by test-retest reliability studies that use language stimuli or that explicitly pertain to the reliability of regions involved in language processing. These show that stable individual participant activity patterns and networks are more variable and distributed than the set of aforementioned ‘language regions’^15–20^. Though there are a number of reasons this might be, a significant amount of the variance can be accounted for by individual differences in task strategies and cognitive style, like the tendency to visualize words^19,21^.

A more specific example derives from the neurobiology of lexical processing. Words activate sensory and motor regions associated with their meaning, including ‘language regions’ and far beyond. The written word ‘telephone’ activates auditory cortex more than words that do not have auditory connotations^22^, ‘red’ activates the visual colour region V4^23^, ‘kick’ activates dorsal motor regions involved in moving the legs^24^ and ‘garlic’ activates olfactory cortex^25^. These activity patterns occur early, within 50-150 ms of word onset, while they are still being read or heard^22,26–28^. This implies that the increased involvement of sensory and motor regions is not simply a post-perceptual process, somehow separable from true ‘language regions’.

These test-retest reliability and word processing examples suggest that ‘language regions’ or ‘the language network’ underrepresent the distributed nature of the neurobiology of language. The difference between the empirical data and popular theoretical models might be explained by the reliance on measures of central tendency. That is, the majority of experiments upon which these models are (presumably) based, average over individuals’ unique though reliable differences in language related activity patterns. This results in ‘language regions’, even if the peaks of any given participant’s blobs are not in those regions^17^. Similarly, the vast majority of studies average over individual words and sentences from many unrelated semantic categories, averaging out their unique distributed patterns, leaving only ‘language regions’.

### Formulaic Language

Individual differences and word processing are only two sources of spatial variability suggesting that the organization of language and the brain is more distributed than suggested by contemporary models. Another possible source of variability, related to both examples, is how well one has learned words and sequences of words. Neuroimaging studies that use multiword or sentence stimuli typically average over word sequences that are more or less formulaic. Formulaic language can be defined as:

> *A sequence, continuous or discontinuous, of words or other meaning elements, which is, or appears to be, prefabricated: that is, stored and retrieved whole from memory at the time of use, rather than being subject to generation or analysis by the language grammar*.^*29*^

Formulaic language is synonymous with or includes many other terms like ‘automatic speech’, ‘chunks’, ‘collocations’, ‘holistic’, ‘fixed, ‘multiword expressions’ ‘noncompositional’, ‘preassembled’, ‘prefabricated’ and ‘ready-made’, to name ten^29,30^. Examples range from explicatives (‘shit’^31^), idioms (‘sort of’^32^), lists (‘one, two, three’), high frequency n-grams (‘the end of the’^33^), to individualised overlearned material (such as ‘I have of late, but wherefore I know not, lost all my mirth’^34^). Formulaic expressions are important for first and second language acquisition^35–37^ and are ubiquitous in everyday usage, comprising a third or more of language^38^. They are processed faster with fewer errors in both children and adults compared to novel words, attesting to their special status^39–44^.

Indeed, when they are not averaged together with less formulaic words, there is evidence that formulaic language processing might have a more distributed pattern than is suggested by popular neurobiological models of language. That is, since the 19th century, it has been known that the ability to produce formulaic speech is often preserved in aphasia, even with severe language impairment and damage to most and sometimes all of ‘the language network’ (as in global aphasia)^45–54^.

Where in the brain are formulaic expressions stored and processed when ‘language regions’ have been destroyed? One possibility is suggested by the relative behavioural and corresponding cortical preservation in aphasia and Alzheimer’s disease. Lesion locations strongly associated with language do not typically include sensorimotor regions around the central sulcus, i.e., primary motor and somatosensory regions^55–62^. Similar to aphasia, people with Alzheimer’s disease also produce more formulaic language than controls^63–66^ and the amount of formulaic language produced predicts disease progression^65^. In addition to formulaic expression production, individuals with Alzheimer’s disease retain other motor functionality. This corresponds to the relative degradation of temporal cortices and sparing of regions and connectivity around the central sulcus^67–73^. In contrast, individuals with Parkinson’s disease produce less formulaic language^63,66,74,75^, have more motor problems and this corresponds to a breakdown of cortical/subcortical sensorimotor networks^76–78^.

### Speech Perception

This work suggests that preserved formulaic language production in aphasia and Alzheimer’s disease is more reliant on sensorimotor regions and less on typical ‘language regions’ or ‘the language network’. By extension, this implies that, as learning increases and words become more formulaic, they become more reliant on these same sensorimotor regions in healthy individuals. In some models of speech perception, it has been suggested that regions of the brain supporting speech production, including sensorimotor regions, are also variously involved in speech perception^79^. As such, the perception and comprehension of formulaic expressions might also rely more on sensorimotor regions and less on ‘language regions’.

This suggestion is supported by neuroimaging studies of music and speech learning. These collectively show that perception after learning involves more engagement of the production systems used during learning. For example, in both monkeys and humans, learning to play sounds on a keyboard is subsequently associated with specific activation of sensorimotor regions involved in making finger and hand movements when listening to those sounds^80–85^. More generally, musical training improves sound and speech perception and these improvements are typically associated with sensorimotor regions^85–90^. Similarly, learning new speech sounds, novel words and artificial languages are all associated with more sensorimotor activity during perception following learning and consolidation^79,91–99^.

What mechanistic account makes sense of ‘hearing’ more with sensorimotor cortices and less with ‘language regions’? All listeners are faced with the problem of achieving perceptual constancy during speech perception. This is because there is variance in acoustic patterns both across and within talkers and no, as of yet, discoverable mapping between these patterns and speech categories (i.e., phonemes, syllables, etc.). We and others have argued that this difficulty might be solved if, during perception, the brain makes use of the contextual information that accompanies speech and the capacity of motor systems to predict the sensory consequences of movements (an ability important for motor learning and adjusting movements in real time, known as ‘efference copy’)^12,79,100,101^.

For example, hearing ‘She was tired of her life and felt ready for a …’ preactivates ‘change’. This is sequenced by motor regions involved in speech production as if it were to be spoken. Through efference copy from sensorimotor regions, the motor pattern for producing ‘ch’ activates acoustic patterns for ‘ch’ in auditory cortices. If there is an overlap with incoming acoustic information, interpretation is confirmed and further processing of ‘change’ is unnecessary, conserving metabolic resources (relative to a less predictive context). Indeed, there is a large reduction in activity in the entire superior temporal plane in a variety of more predictive contexts during speech perception and language comprehension^101–105^. By this account, the more overleared and formulaic a sequence of words, the earlier the whole sequence becomes predictable. In the example, ‘change’ might be predictable at ‘ready’ before overlearning and ‘tired’ after. Correspondingly, perception will be more supported by sensorimotor regions and much less by ‘language regions’.

### Hypothesis

Based on these speculations, we tested the hypothesis that, as sentences become formulaic through production-based overlearning, there will be a reorganization of the brain regions supporting the perception of those sentences. Behaviourally, we operationalised overlearning as a decrease in reaction times to predict the final word of two sentences, to process the individual words in those sentences and the ability to accurately remember the sentences following two weeks of home production based listening and repetition.

Neurobiologically, reorganization was predicted to correspond to a large increase in activity or new activations of sensorimotor regions. Here and throughout, we define ‘sensorimotor regions’ as pre and primary motor and primary and secondary somatosensory cortices in and adjacent to the central sulcus. These and other regions were expected to support the production of novel sentences. Concomitantly, we predicted a large decrease or inactivity in ‘language regions’ while listening to overlearned sentences. ‘Language regions’ were defined as activation when listening to novel sentences not previously heard or overlearned. Finally, we expected these changes to correspond to a global network reorganization of the brain.

Importantly, we used a natural or so called ‘passive’ design in which participants listened without making meta-linguistic judgements or corresponding button or vocal responses. This is because our hypothesis maintains that sensorimotor regions involved in production support perception after overlearning. If we had included motor responses, there would be no verifiable way to argue that resulting sensorimotor activity was not due to those movements. Figure 1 provides an overview of the study design used to test these predictions.

**Figure 1.**
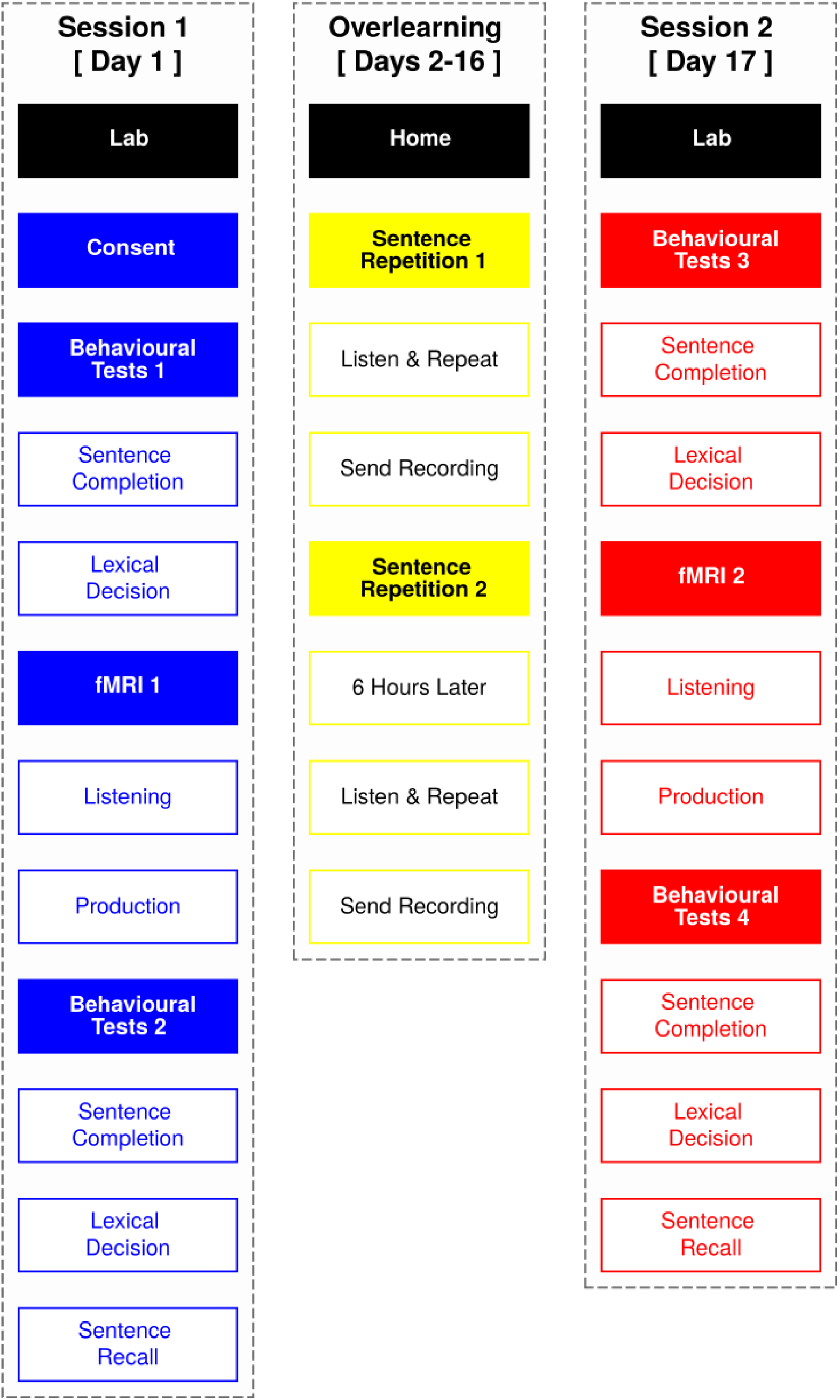
Study overview. We conducted functional magnetic resonance imaging (fMRI) over two sessions. In both sessions, participants ‘passively’ listened to two sentences repeated 60 times each and 60 novel sentences. In a final run, participants produced the 60 novel sentences. The sessions were separated by 15 days. During this time period, participants trained at home by producing the two repeated sentences from fMRI 30 times each, twice a day. To assess learning, participants performed sentence completion, lexical decision and sentence recall tasks before and after training.

## Methods

### Participants

Twelve native British English speakers participated (6 females; 21-25 years old; M = 23.17; SD = 1.41). All but one was monolingual. Participants were right-handed as determined by the Edinburgh Handedness Inventory^106^. All had unimpaired hearing and (corrected) vision. None had any contraindication for MRI, history of psychiatric or neurological disorder or language-related learning disabilities. All participants gave informed consent and the study was approved by the University College London ethics committee.

### Procedure

The experiment lasted 17 days, including two testing days, each with three different behavioural tasks (completed multiple times) and fMRI to assess overlearning of two sentences (Figure 1). On the first day, participants performed a sentence completion and lexical decision task on a desktop computer in a noise attenuated testing room, using headphones. Both tasks included the words from the two sentences that the participants would overlearn over the subsequent 15 days.

Following these tasks, participants were escorted to the scanning suite. There they chose comfortable earbud sizes for noise-attenuating headphones. After being instructed, they were put in the head-coil with pillows under and on the sides of their head and (if desired) under the knees for comfort and to reduce movement during scanning. Once in place, participants chose an optimal stimulus volume by determining a level that was loud but comfortable. Once scanning began, participants’ first fMRI task was to ‘passively’ listen to the two ‘overlearned’ sentences multiple times and previously unheard or ‘novel’ sentences. Participants’ last fMRI task was to listen to and repeat some of the novel sentences that they had earlier heard in the scanner. Finally, we acquired high-resolution anatomical scans. After scanning, participants were returned to the testing room where they did the sentence completion and lexical decision tasks a second time and a sentence recall task to assess learning over the first day.

Over the next 15 days, participants trained at home by listening to and repeating the two overlearned sentences they heard during fMRI. They did this twice a day, sending us recordings of their productions when they were done. Participants returned on the final (17th) day and performed the sentence completion, lexical decision and sentence recall behavioural tasks, fMRI passive listening and speech production tasks and anatomical scans as on day one, in the same order. When these were complete, participants were given £7.5 per hour for behavioural testing and £10 per hour for scanning to compensate for their time and sent home.

### Stimuli

Participants were divided into three groups. Each group of four participants overlearned two different sentences. Three pairs of sentences were used to help assure that results are generalizable. Specifically, a male talker recorded the 498 sentences from the Supplemental Materials of Block and Baldwin^107^. Sentences were edited in Praat (http://www.fon.hum.uva.nl/praat/) to be 2.5 seconds, with the final word lasting 500 ms (see next ‘Sentence Completion’ section for a rationale). Four raters judged whether the sentences were appropriate for British English listeners (e.g., they do not discuss American football or ‘pants’) and sounded natural (i.e., they were not sped up or slowed down). The latter was done using a Likert scale from one (not natural) to 10 (very natural). Inappropriate sentences, those with a naturalness rating less than six and sentences with proper nouns were discarded. From the remaining sentences, three sets of two sentences were created that were matched on number of words, cloze probability (i.e., the probability of the final word completing the preceding words)^107^, complexity and average word frequency as determined by Subtlex-UK^108^ (Table 1). From the remaining sentences, 60 high cloze probability sentences were selected to be used in behavioural tasks and during scanning.

**Table 1.** Overlearned sentence information. All sentences were 2.5 seconds long and the final word was 500 ms. Cloze probability from Block and Baldwin (2010)^107^. Complexity and word frequency are from Subtlex-UK^108^.

| Set | Sentence | Words | Cloze Probability | Complexity | Average Frequency |
| --- | --- | --- | --- | --- | --- |
| 1 | Instead of dressing, I prefer vinegar and oil. | 8 | 0.76 | 1.2 | 7716.87 |
|  | At the pub, he ordered another mug of beer. | 9 | 0.87 | 1.2 | 8645.02 |
| 2 | She was tired of her life and felt ready for a change. | 12 | 0.83 | 1.3 | 7134.33 |
|  | The jury found him innocent and set him free. | 9 | 0.85 | 1.3 | 7996.94 |
| 3 | They went to the video store to rent a movie. | 10 | 0.54 | 1.4 | 13093.97 |
|  | She followed the recipe correctly to cook the meal. | 9 | 0.56 | 1.4 | 13651.17 |

### Behavioural Tasks

Three behavioural tasks were used to assess overlearning of the two sentences assigned to each participant. Sentence completion and lexical decision tasks were completed both before and after the two fMRI sessions. The sentence recall task was done only after each imaging session.

### Sentence Completion

Participants listened to two sentences that were to be overlearned (sessions one and two) or had been overlearned (sessions three and four) and 30 novel sentences. In each case, the final word in the sentence was removed. Participants were asked to press a button as soon as they believed they knew the final word. The sentence stopped playing when the button was pressed and participants then typed in the final word as quickly as possible. Reaction times for the task were measured from the start of sentence playback until participants finished entering the predicted final word. On a per participant basis, reaction times greater than four standard deviations from the mean reaction were not included in the analysis.

### Lexical Decision

Participants had to judge the lexical status (word or not word) of overlearned words, words similar to overlearned words, words dissimilar to overlearned words and nonwords. Thirty overlearned words were extracted from the overlearned sentences used in the study. Eighteen similar words were selected by searching through the remaining novel sentences and finding words that had a Levenshtein distance of two or less from the overlearned words extracted from the overlearned sentences. Levenshtein distance is the minimum number of single character alterations required to change one word into another^109^. Sixteen dissimilar words were also identified. These had a Levenshtein distance of six or more and also had a frequency that differed by ±10% from the overlearned words. Sixty-four unintelligible but speech-like nonwords were created from the selected (i.e., overlearned, similar, dissimilar) words using a local time reversal script in Praat with 150 ms steps^110^.

Each participant heard only 21 words (nine overlearned words taken from the two sentences they repeated, six similar words and six dissimilar words) and 21 nonwords (reversed versions of the overlearned, similar and dissimilar words they heard). After hearing each word they indicated if they heard a word or a non-word with a button-press as quickly and accurately as possible. Reaction times were measured from the start of playback to the moment a response was indicated. On a per participant basis, reaction times greater than four standard deviations from the mean reaction were not included in the analysis. We then examined how reaction times changed between the two scanning sessions.

### Sentence Recall

After scanning sessions, participants were asked to type in up to 10 sentences that they remembered hearing in the scanner. The sentences they recalled (e.g., ‘He was a lousy cook and ordered out’) were matched (based on semantics and word use) to the sentences that were actually played in the scanner (e.g., ‘He eats out because he is a lousy cook’). The cosine similarity was then found between the two sentences to provide a measure of recall accuracy. The cosine similarity is the cosine of the angle between two normalized vectors. In this case, the two sentences were transformed into vectors of word counts and the cosine similarity was found for these word count vectors. Cosine similarity takes a value between zero and one, where a value of zero means that the sentences share no words and a value of one means that the sentences are identical in terms of words used.

### Sentence Overlearning

Commencing the day after the first fMRI session, participants overlearned two sentences through repetition at home, twice a day. Each day, participants listened to a prerecorded set of their two sentences, each repeated 30 times in a random order, lasting five minutes and 32 seconds. There was a three second gap between sentences during which the participant repeated the sentence out loud. The participants did this task again at least six hours later using another randomised prerecorded file of the same two sentences. All 30 prerecorded files had a different randomisation. To verify home learning took place, participants recorded themselves listening to and repeating their sentences using Audacity (http://www.audacityteam.org) and immediately shared the recording to a cloud storage folder. In total, participants listened and repeated their two sentences 1,800 times for two hours and 46 minutes. This was verified by checking recordings.

### fMRI

#### Task

A slow random event related design was used to compare how the response to overlearned sentences changed from fMRI session one to two. In each scanning session, audio stimuli were presented during six listening runs and a speech production run. Each listening run lasted six minutes and 53 seconds. Across these, participants heard two sentences repeated 60 times each and 60 novel sentences that were each only heard once, all in a randomised order. Stimuli were presented in a jittered manner such that, following each 2.5 second sentence, there was a minimum of 10 seconds of silence and a mean of 10.675 seconds (SD = 0.93) and a maximum of 15 seconds of silence. The speech production run always followed the six listening runs and lasted eight minutes and 10 seconds. In this run, participants listened to 30 of the novel sentences they had just heard and were asked to repeat each sentence as soon as it finished. After each 2.5 second sentence and allowing for another 2.5 second period to produce the sentence, there was, again, a minimum of 10 seconds of silence with a mean of 10.69 seconds (SD = 1.34) and a maximum of 15.625 seconds of silence. All listening and production runs included 10 seconds of silence at the start to allow magnetization to reach a steady state. Sessions one and two were the same, with the exception that in session two they produced the other 30 sentences not produced in session one. Participant engagement was monitored with a camera over one eye.

Functional and anatomical images were acquired on a 1.5T Siemens MAGNETOM Avanto with a 32 channel head coil (Siemens Healthcare, Erlangen, Germany). We used multiband EPI^111,112^ (TR = 700 ms, TE = 54.8 ms, flip angle of 75°, 28 slices, resolution = 3 x 3 x 4 mm), with 4x multiband factor and no in-plane acceleration. Slices were manually obliqued to include the entire brain. The six listening EPI runs were each 590 volumes/TRs and the speech production run had 700 volumes/TRs. Two six minute T1-weighted high-resolution MPRAGE anatomical MRI scans followed the functional scans (TR = 2.73 s, TE = 3.57 ms, 176 sagittal slices, resolution = 1.0 mm^3^). Imaging parameters were the same for both fMRI sessions (thus resulting in 12 listening, two production runs and four anatomical scans).

### Preprocessing

Unless otherwise noted, the AFNI software suite was used for preprocessing and analyses (http://afni.nimh.nih.gov/afni)^113^ Individual AFNI programmes are indicated parenthetically in subsequent descriptions.

The four anatomical/structural MRI scans were corrected for image intensity non-uniformity (‘*3dUniformize*’) and deskulled using *ROBEX*^*114*^. Within each session, the second anatomical image was aligned to the first and they were averaged. Then the resulting session one and two anatomical images were aligned and averaged. This was done using a procedure to reduce bias by moving both anatomical images, so that both are interpolated some amount rather than one session receiving all the interpolation (see https://sscc.nimh.nih.gov/sscc/dglen/alignmentacross2sessions).

The resulting anatomical image was nonlinearly aligned (using ‘*auto_warp*.*py*’) to the MNI N27 template brain, an average of 27 anatomical scans from a single participant (‘Colin’)^115^. The anatomical scan was inflated and registered with Freesurfer software using ‘*recon-all*’ and default parameters (version 6.0, http://www.freesurfer.net)^116,117^. This included automatic parcellations of the anatomical image. These were used to create white matter and ventricle (i.e., cerebral spinal fluid containing) regions of interest that were used as noise regressors. Automatic parcellation was also used to generate 167 regions of interest for network analyses (i.e., using the ‘Destrieux Atlas’)^117^.

The first 10 TRs (7 seconds) were removed from the fMRI timeseries before they were corrected for slice-timing differences (‘*3dTshift*’) and despiked (‘*3dDespike*’). Next, volume registration was done by aligning each timepoint to the mean of run four (‘*3dvolreg*’). The functional data were then aligned to the anatomical images (‘*align_epi_anat*.*py*’). This used the less biased procedure described for anatomical alignment, moving functional data from both sessions so that no one session received all the interpolation. Finally, the volume-registered and anatomically-aligned functional data were (nonlinearly) aligned to the MNI template brain (‘*3dNwarpApply*’).

Next, we created two sets of timeseries. The first, to be used in the deconvolution analysis described below, involved only normalising each run to have a sum of squares of one (‘*3dTproject*’). The second set of time series were normalised and detrended using Legendre polynomials whose degree varied with run lengths (following the AFNI recommended formula of [number of timepoints * TR]/150). These were then submitted to spatial independent component analysis (ICA) to detect and remove artifacts. Specifically, we concatenated the normalised and detrended listening and production timeseries from both sessions separately (‘*3dTcat*’). We did ICA on the resulting listening timeseries with 300 dimensions and on the production timeseries with 100 dimensions using ‘*melodic*’ (version 3.14) from FSL^118^. Next, we labelled and removed artifacts from the timeseries, following recommendations from an existing guide for manual classification^119^. One of two trained authors went through all components and associated timecourses, labelling the components as ‘good’, ‘maybe’ or ‘artifact’. Our strategy was to preserve signal by not removing components classified as ‘maybe’.

Finally, we made a third timeseries using the concatenated listening runs for the regional homogeneity analysis described below. Specifically, the timeseries were normalised to have a sum of squares of one and detrended (‘*3dTproject*’) with the following regressors: 1) Legendre polynomials whose degree varied with run lengths (following the previously described formula); 2) Six demeaned motion regressors from the volume registration (roll, pitch, yaw and changes in the inferior/superior, left/right and anterior/posterior directions); 3) A demeaned white matter activity regressor from the averaged timeseries in white matter regions; 4) A demeaned cerebrospinal fluid regressor from the averaged timeseries activity in ventricular regions; and 5) the ICA artifact component timecources.

### Individual Deconvolutions

After preprocessing, two individual participant deconvolutions were conducted to get an estimation of the system impulse response function for the 1) overlearned and novel sentences from the listening runs and the 2) produced novel sentences from production run from both fMRI sessions (*‘3dDeconvolve*’)^120,121^. In the first deconvolution, regressors of interest included one each for overlearned sentence one in session one, overlearned sentence two in session one, novel sentences in session one, overlearned sentence one in session two, overlearned sentence two in session two and novel sentences in session two. For each of these, the hemodynamic response was estimated using a cubic spline basis function that covered an 18 second period after each stimulus onset, using 20 tent functions to generate the impulse response function for every voxel. In a second deconvolution, regressors of interest included speech production of novel sentences in session one and speech production of novel sentences in session two. Again, the cubic spline basis function was used with the difference that the period covered was 20 seconds (to account for the extra time involved in producing the sentences), using 21 tent functions.

In both deconvolutions, regressors of noninterest included an automatically estimated number of polynomials (again following the [number of timepoints * TR]/150 formula), six motions regressors from the volume registration step (roll, pitch, yaw and changes in the inferior/superior, left/right and anterior/posterior directions), two regressors from the average timeseries in the white matter and ventricles and all timecourses from ICA components labelled as artifacts.

### Novel LMM

Following individual participant deconvolutions, the resulting impulse response functions were spatially smoothed to achieve a level of smoothness of six mm full-width half maximum, regardless of the smoothness it had on input (‘*3dBlurToFWHM*’^122^). These were then used in four linear mixed-effects models (LMMs; ‘*3dLME*’)^123^.

First, we did a novel sentence listening LMM. Factors were session (one and two) and timepoint (0-25). This allowed us to test the prediction that learning was specific to overlearned sentences. Though we did not expect differences in activity for the novel sentences across sessions, participants might have learned something about the talker’s voice as the same talker made all sentences used in the study. Time was included as a factor to compare to the overlearning LMM (see next paragraph) though, again, we did not expect that there would be differences in the shapes of the hemodynamic response between sessions one and two. There were 26 timepoints because our TR was 700 ms and there are, therefore, 26 TRs covering the 18 second period from the deconvolution. Finally, collapsing over session and time, the novel LMM served to identify ‘language regions’ which were expected to encompass the bilateral inferior frontal gyrus and superior temporal plane.

### Overlearning LMM

Second, we conducted an overlearning sentence listening LMM with sentence (overlearned sentence one and two), session (one and two) and timepoint (0-25) as factors. This allowed us to understand the effect of learning on overlearned sentence listening between sessions one and two. Time is included as a factor because we expect differences in the shapes of the hemodynamic response across sessions though we did not make a priori predictions about the direction of those differences in individual brain regions. We also visualised the timepoints from the results to better understand if responses for overlearned sentences are a simple redistribution of activity (i.e., a relative modulation of activity from session one to two) or a reorganization of brain responses (i.e., activity in regions in session one or two that was not previously present).

### Overlearning-Novel LMM

Third, a followup analysis was conducted by subtracting the novel from the overlearned impulse response functions at each timepoint in each participant. We then ran a LMM with the same factors as the overlearning sentence listening LMM (i.e., session*sentence*timepoint). This allowed us to formally test whether overlearned sentences produced significantly more activity than novel sentences in sensorimotor regions and less activity than novel sentences in ‘language regions’. We present the results of each session separately so that the direction of effect can be interpreted.

### Production LMM

Finally, we did a novel speech production LMM with session (one and two) and timepoint (0-28) as factors. This analysis was used to demonstrate regions involved in producing the novel sentences that participants heard during both sessions.

To correct for multiple comparisons in all LMMs, we used a multi-threshold approach rather than choosing an arbitrary p-value at the individual voxel level as is customary. In particular, we used a cluster simulation method to estimate the probability of noise-only clusters using the spatial autocorrelation function from the residuals in each LMM (‘*3dFWHMx*’ and ‘*3dClustSim*’). This resulted in the cluster sizes to achieve a corrected alpha value of 0.01 at nine different p-values (i.e., 0.05, 0.02, 0.01, 0.005, 0.002, 0.001, 0.0005, 0.0002 and 0.0001). We thresholded each map at the corresponding z-value for each of these nine p-values and associated cluster size. We then combined the resulting maps, leaving each voxel with its original z-value.

### Regional Homogeneity

The described deconvolution approach uses a linear model to derive an estimate of the hemodynamic response from multiple stimulus presentations. We reasoned that a strong case for the hypothesis could be made if a similar set of results were obtained using a more model-free approach across the whole timeseries. To do this, we used regional homogeneity which calculates the Kendall’s coefficient of a concordance for each voxel within a neighbourhood of voxels (‘*3dReHo*’)^124,125^. We chose this particular approach because it is also a measure of local interactions, synchronisation and connectivity^125^. Corresponding to our hypothesis, we expected an increase in local connectivity in sensorimotor regions and a decrease in superior temporal plane and inferior frontal regions after overlearning.

To do this analysis, we first constructed three timeseries that theoretically reflect only timepoints for processing overlearned sentences from session one, overlearned sentences from sessions two or novel sentences from both sessions. Specifically, we modelled expected hemodynamic responses by convolving stimulus onsets with a canonical hemodynamic response function (using ‘WAV’, a.k.a the ‘Cox special’ from ‘*waver*’). We then cut up the third timeseries described in ‘Preprocessing’ above by taking the relevant timepoints under the canonical response starting from the timepoint that the response starts to rise (a delay of 2.1 seconds or three TRs) and ending when the response returns to baseline (*‘3dTcat’*). We removed any timepoints that overlapped in any of the timeseries. Because this was a slow event related design with jitter, this amounted to only 7.02% of the data that was about equally distributed across the three sentence timeseries.

We then did the regional homogeneity analysis for each of the three resulting timeseries using a radius of 2.3 which equals 57 voxels (we also tested 2.0 or 33 voxels and 2.9 or 93 voxels and it makes little difference to the results). The resulting maps were then blurred to achieve a level of smoothness of six mm full-width half maximum (‘*3dBlurToFWHM’*). After this, we conducted three group paired t-tests to compare 1) the overlearned sentences from session one to those from session two; 2) the novel sentences from sessions one and two to the overlearned sentences from session one; and 3) the novel sentences from sessions one and two to the overlearned sentences from session two (‘*3dttest*++’).

We saved the results as z-scores and used the residuals from the t-tests to do the same multi-thresholding procedure described for the LMM analyses to correct for multiple comparisons. We note that *3dttest*++ has a built-in and more sophisticated equitable thresholding procedure (called ETAC)^126^. However, at the time of analysis, there was no obvious way to use this approach with, e.g., the LMM results. As such, we opted to use our described approach for consistency across analysis. That said, we looked at the results using the ETAC approach for comparison and they look similar to our multi-thresholding approach.

### Network

To do network analyses, we first concatenated the two overlearned sentence impulse response functions from the deconvolution for the two sessions separately. We blurred these as in previous analyses to a level of smoothness of six mm full-width half maximum (‘*3dBlurToFWHM’*). Using the Freesurfer ‘Destrieux Atlas’ parcellation, 167 regions of interest were extracted from these timeseries for each participant. Pairwise Pearson’s correlations were used to build two unweighted, undirected adjacency matrices for each participant, one each for sessions one and two overlearned sentences. Absolute thresholding of r = 0.1 was applied to correlation values, in order to build adjacency matrices for group comparisons^127,128^.

To test whether global network reorganization took place after overlearning, the distance measures ‘edit distance’ and ‘Deltacon’ were calculated. Edit distance computes additions or deletions of connections between two graphs^129^. The edit distance matrix was defined as:

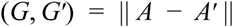

Where A and A’ are the adjacency matrices for graphs G (session one) and G’ (session two) respectively, and δ is the pairwise edit distance^129^. Since session one and two shared node identity, this pairwise application was applicable. Change in connectivity was calculated as the ratio between the total number of lost and gained connections in the edit distance matrix and the total number of connections in both session one and two adjacency matrices for a single participant. A one-sample t-tests across participants was performed on the change in connectivity values to determine if they differed from the null, i.e, no change in connectivity.

In order to describe and visualise connections involved in edit distance differences from session one to two, a two-way chi-square test was performed on each region of interest pair across participants. Lost connections were defined as those whose connectivity in session two was lower while observed connectivity in session one was higher than expected. Gained connections were defined as the converse. We set a threshold at p < .01 to afford some protection for multiple comparisons.

Though the edit distance matrix is simple to compute, it suffers from limitations. It only determines specific connection changes, but it does not interpret the change in the context of the rest of the network and its neighbours. Moreover, it does not differentiate between network densities: if a connection is lost in a very sparse network the result would be a large disruption, but if a connection is lost in a highly dense network the outcome on the global scale will be minimal^130^. For these reasons we also calculated Deltacon, a more robust similarity measure that determines the level of isomorphism between two networks with node correspondence, using Matusita’s distance^130^. We compared the results using a one-sample t-test across participants on the deltacon dissimilarity value (i.e. 1-Deltacon).

To further examine possible changes between session one and two, we explored a number of other global network measures. These included efficiency (average inverse shortest path length)^131^, diffusion, density and flow using the Brain Connectivity Toolbox in MATLAB^132^. Density measures how ‘connected’ a network is, diffusion measures how quickly information can get from point A to B and flow measures how centralised a network is for transfer of information^132^. We used a two-sample t-tests for these global measures, comparing sessions one and two. Finally, we computed two measures of local connectivity, centrality and community partitioning. Centrality (degree, eigenvector, closeness and betweenness) measures the importance of a node in a network, whilst community detection partitions the network into distinct sub-components or modules^132^.

## Results

### Behavioural Tasks

We hypothesized that participants would show behavioural markers of overlearning for the sentences they repeated at home. We assessed this with three behavioural tasks (Figure 2).

**Figure 2.**
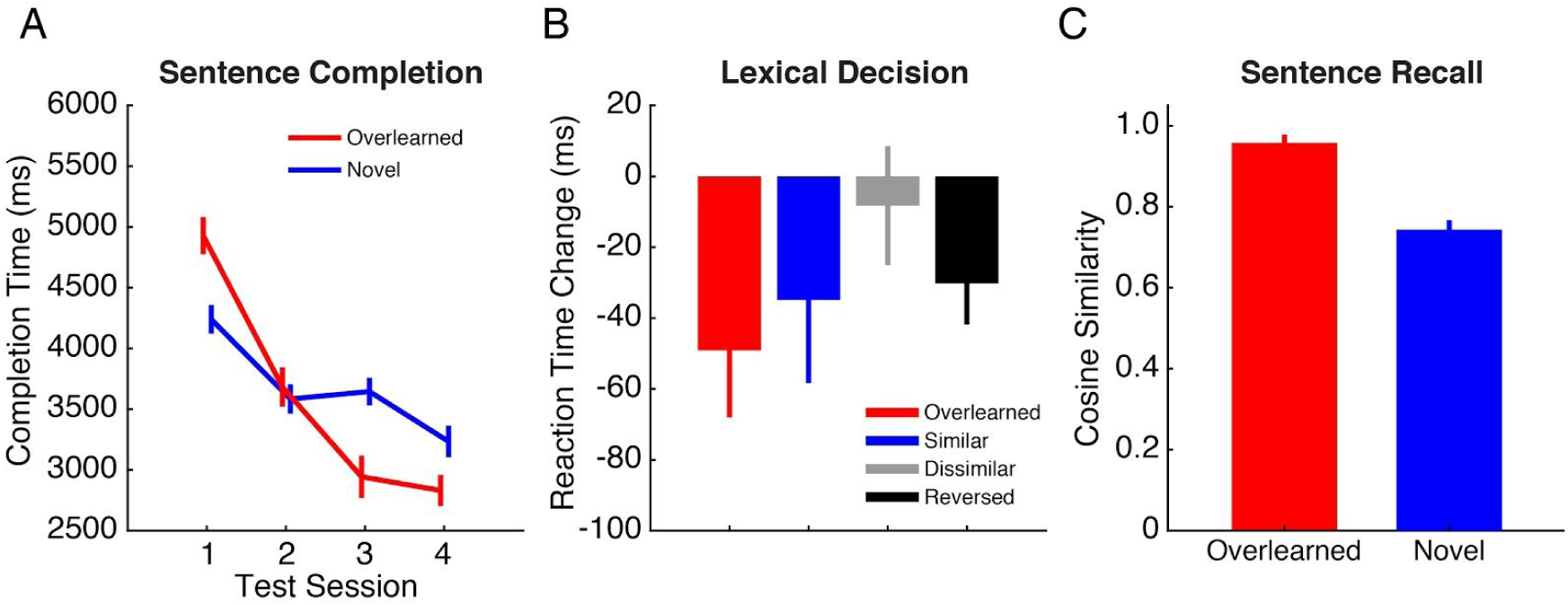
Behavioural measures of overlearning. A) For each testing session (1-4), sentence completion is the length of time (in ms) that it took participants to type in the final word of overlearned (red line) and novel sentences (blue line). Completion times decreased following overlearning. B) Lexical decisions (word / not word) were made for words drawn from the overlearned sentences (red bar), similar words (blue bar), dissimilar words (grey bar) and reversed words (black bar). Change in lexical decision time reflects decision times (in ms) measured in sessions one and two (pre overlearning) versus decision times measured in sessions three and four (post overlearning). C) Participants recalled sentences they heard during scanning. The accuracy of sentence recall was measured as the cosine similarity between recalled and heard sentences. Participants were more likely to recall sentences they had overlearned (red bar) versus novel sentences (blue bar).

### Sentence Completion

Figure 2A shows sentence completion times (in which participants completed the final word of a sentence) across testing sessions for both overlearned and novel sentences. There was an interaction between completion time measured at each session and whether the sentence was overlearned or not (F(3,33)=22.28, p < .001). For overlearned sentences, a decrease in sentence completion times was observed following training (session two versus session 3; t(11) = 3.90, p < .01). A similar decrease was not observed for novel sentences (t(11) = -1.04, p = .32). By the end of the study, participants completed the final word in the overlearned sentences 2.10 seconds faster then they did at the start of the experiment. By comparison, the time it took to complete the final word in novel sentences improved by 1.0 seconds.

### Lexical Decision

Figure 2B shows changes in lexical decision times following overlearning (testing sessions one and two versus sessions three and four), for words drawn from overlearned sentences (red bar), words similar to overlearned words (blue bar), words dissimilar to overlearned words (grey bar) and nonwords (black bar). Changes in lexical decision time between the four word types were not statistically significantly different (F(3,33) = 2.32, p = .094). However, words drawn from overlearned sentences were identified faster following overlearning than they were prior to training (t(11) = 2.59, p = .025). A comparable result was not observed in the cases of similar (t(11) = 1.49, p =.16) and dissimilar words (t(11) = .49, p = .63).

### Sentence Recall

Finally, Figure 2C gives a measure of sentence recall (i.e., memory) for overlearned sentences versus novel sentences following overlearning. Participants recalled overlearned sentences heard during scanning with significantly greater accuracy than novel sentences heard during scanning (t(11) = 6.54, p < .001). In fact, half of the participants recalled the overlearned sentences verbatim. By comparison, none of the participants recalled the novel sentences verbatim even though they had just heard them during scanning.

### fMRI

#### Novel LMM

We expected that learning would be specific to overlearned sentences. To help determine if this was the case, we conducted a novel sentence listening linear mixed-effects model (LMM) with session (one and two) and timepoint (0-25) as factors. There were no discernible effects of session and no session by time interaction. Nonetheless, we used general linear tests (GLTs) to directly contrast novel sentences for sessions one and two at all timepoints in a manner used in subsequent analyses (see Figure 3). Collapsed over all 26 timepoints, there were a small number of differences from session one to two (Figure S2). This included seven clusters with more than 20 voxels, with decreases in activity in the cerebellum (x/y/z = 31/-84/-41; 890 voxels), thalamus (x/y/z =-5/-15/7; 42 voxels) and lingual gyrus (x/y/z =-29/-48/-2; 25 voxels) and increases in the left dorsal postcentral gyrus (x/y/z = -41/-39/67; 657 voxels), superior frontal gyrus (x/y/z =-1/32/36; 161 voxels), right superior parietal lobule (x/y/z =19/-57/70; 99 voxels) and right middle anterior cingulate gyrus (x/y/z =4/18/25; 20 voxels). We calculated the GLTs for novel sentences for sessions one and two independently and used these for analysis of the impulse response function (described in the next section; see Figure 4). We also calculated the GLTs for novel sentences, collapsing over sessions one and two and all 26 timepoints and used this as a guide to ‘language regions’ (see Figures 3 and 5).

**Figure 3.**
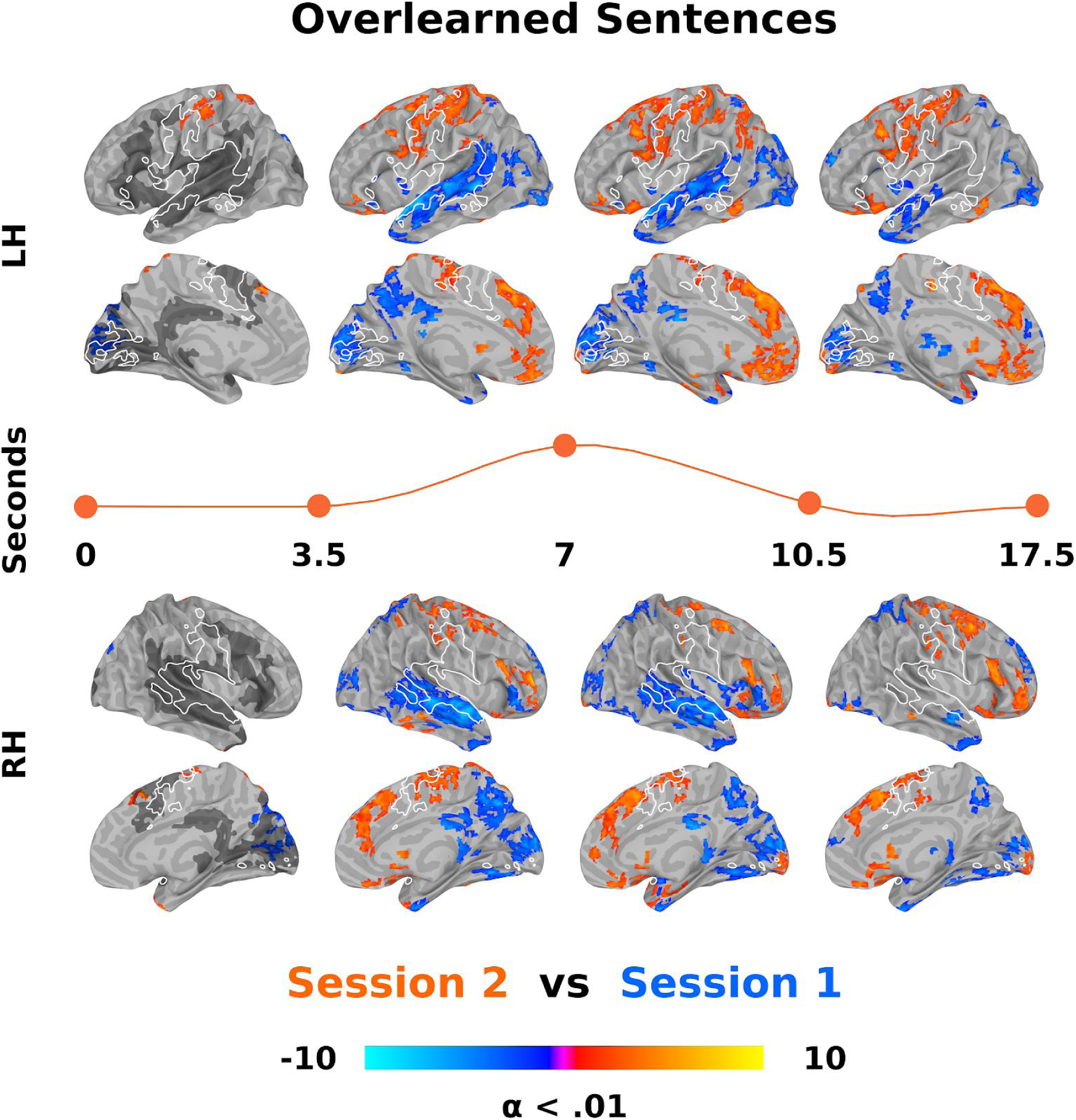
Direct contrast of overlearned sentence listening from session one and two across time. General linear tests (GLTs) between session one and two overlearned sentences were done at each of 26 timepoints in the estimated impulse response functions following a linear mixed-effects model (LMM). For ease of visualisation, these contrasts are collapsed into four time bins between the indicated seconds. The orange line separating the left (LH) and right (RH) hemisphere surfaces roughly represents the relative position in a ‘canonical’ hemodynamic response function. Listening to sentences after overlearning results in an increase in activity in sensorimotor regions centred around the central sulcus (among other regions, reds) and a reduction of activity in the superior temporal plane (blues). Increased sensorimotor activity overlaps with producing novel sentences as determined by a separate LMM (white outline). Note that the sensorimotor regions are first engaged after overlearning in the delay period before the canonical response rises. Decreased superior temporal plane activity overlaps with listening to novel sentences as determined by another LMM (dark grey, presented on only the far left column though it represents all 26 timepoints). Each timepoint was cluster size corrected for multiple comparisons at alpha (α) < .01. The colour bar represents z-scores.

**Figure 4.**
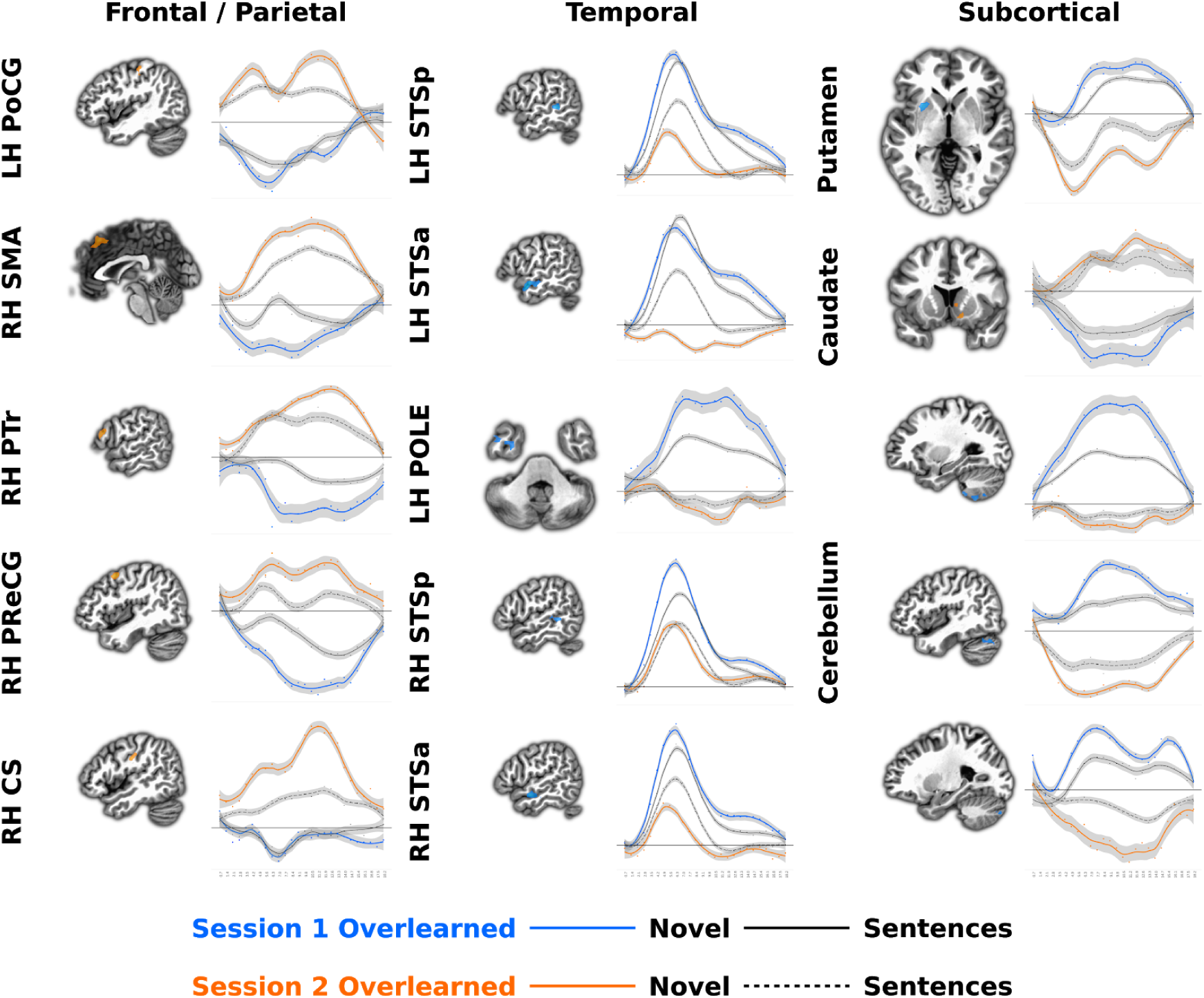
Estimated impulse response functions for overlearned and novel sentence listening in different regions. The direct contrast of overlearned sentences from sessions one and two was thresholded at z = 10 (p < 1.52 x 10^−23^), resulting in 19 clusters of activity. Fifteen of these, five each from frontal / parietal, temporal and subcortical regions were selected for display. The impulse response function from the voxels in each of these clusters from the overlearning linear mixed-effects model (LMM) were averaged and plotted for each session. Also plotted are the averaged novel sentence listening LMM impulse response functions for comparison. Note that in most cases, the change in the impulse response functions before or after learning are from a state of below baseline or inactivity. Abbreviations: CS = central sulcus; LH = left hemisphere; PoCG = postcentral gyrus; POLE = temporal pole; PreCG = precentral gyrus; PTr = pars triangularis of the inferior frontal gyrus; RH = right hemisphere; SMA = supplementary motor area; STSa = anterior superior temporal sulcus; and STSp = posterior superior temporal sulcus.

**Figure 5.**
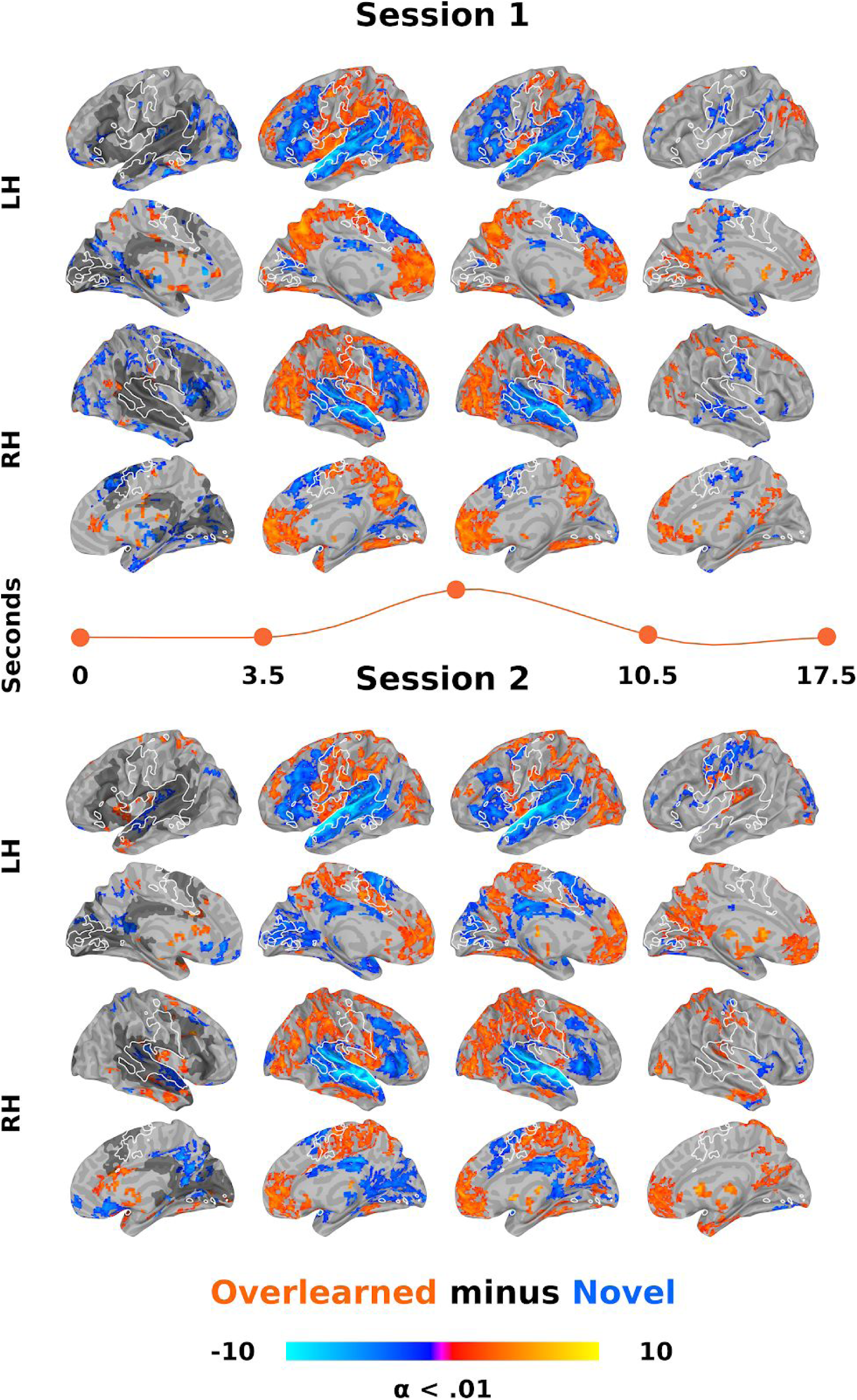
Direct contrast of overlearned minus novel sentence listening for session one and two across time. General linear tests (GLTs) for session one (top) and two (bottom) for the subtraction of novel from overlearned sentences, done at each of 26 timepoints in the estimated impulse response functions. In both sessions, listening to overlearned sentences results in significantly more activity in sensorimotor regions (reds) also involved in producing speech (white outline). Conversely, overlearned sentences result in less activity in the superior temporal plane and inferior frontal gyrus (blues), specifically in regions that are involved in processing novel sentences (dark grey, left hand column). Everything else is as in Figure 3.

### Overlearning LMM

To test the hypothesis that sentence listening after overlearning involves more sensorimotor and less activity in ‘language regions’, a LMM was done with session (one and two), sentence (overleaning sentences one and two) and timepoint (0-25) as factors. There were main effects of session and time that encompassed most of the brain at a corrected threshold. In contrast, the main effect of sentence at p < .01 corrected resulted in only four clusters in the primary visual cortex (x/y/z = -11/-102/4; 615 voxels), middle occipital cortex around motion area MT+ (x/y/z = -47/-72/7; 123 voxels), superior parietal cortex (x/y/z = 28/-51/61; 121 voxels) and middle frontal gyrus (x/y/z = 43/33/34; 117 voxels). Given this relatively small amount of activity and that these regions are mostly centred around visual cortices, suggesting minor differences in sensory semantic properties of the overlearned sentences (recall that there were three sets of sentences), we collapsed over sentences one and two for all subsequent analyses.

Compared to baseline and independent of time, GLTs for sentences in sessions one and two both involve processing in the superior temporal plane. However there was an increase in processing in sensorimotor cortices and a large reduction in the spread of activity in the superior temporal plane in session two (Figure S1A and S1B). A direct contrast reveals that most of the brain differs at corrected threshold and this confirms a bilateral increase in sensorimotor regions and a decrease in the superior temporal plane (Figure S1C). Subcortically, the hippocampus, caudate and dorsal cerebellum increase while the thalamus, putamen and ventral cerebellum decrease from sessions one to two. These results are present even at voxel-wise corrected threshold of p < 1 x 10^−10^.

There was an interaction between session and time for overlearned sentences that involved many of these same regions, suggesting that the timing of activity also changes. To visualize changes, the GLTs for the contrast of sessions one and two at each timepoint are presented (Figure 3). To make visualisation easier, we created four time bins and illustrated approximately when activity occurs with regard to a ‘canonical’ hemodynamic response function (Figure 3, middle line). The previously described results can be seen, with an increase in sensorimotor and a decrease in superior temporal plane regions in session two. Also apparent is that processing began earlier in sensorimotor regions in session two (Figure 3, first column). This includes activity in primary motor and somatosensory cortex in the central sulcus (x/y/z = -44/-24/55; 376 voxels) and the supplementary motor area (x/y/z = 4/21/49; 40 voxels) and no subcortical structures. Differences in subcortical activity began in the second time bin and is as described above (i.e., an increase in activity in the hippocampus, caudate and dorsal cerebellum and a decrease in the thalamus, putamen and ventral cerebellum). Using the separately conducted novel sentence listening and production LMMs as a guide, it appears that activity for overlearned sentences is increased in sensorimotor regions involved in producing speech (Figure 3, white outline) and reduced in regions active during novel sentence listening (Figure 3, dark grey).

Next, we further explored overlearning LMM results by visualising session one and two timecourses for overlearned sentences. This allows us to better understand if responses for overlearned sentences are a simple redistribution or a reorganization of brain responses. Specifically, we thresholded the previously described GLT contrasting overlearned sentences between sessions one and two (see Figure S1C) at an arbitrary high z-value of 10. This resulted in 19 small clusters and we selected 15 of these for presentation, five frontal or parietal, five superior temporal and five subcortical regions. We averaged and plotted the coefficients in each of these regions for all timepoints. For comparison, we also plot timepoints for the separately conducted novel sentence listening LMM (see prior section). Overlearned sentence responses in frontal and parietal, including sensorimotor regions, all showed a new response in session two from something resembling a below baseline or lack of activity in session one (Figure 4, left). In contrast, overlearned sentence responses in superior temporal regions all showed a reduction in, lack of or below baseline response in session two from a state of heightened activity in session one (Figure 4, middle). Finally, subcortical regions showed a similar set of patterns, with an increase in the caudate and a decrease in the cerebellum and putamen for overlearned sentences from session one to two (Figure 4, right).

### Overlearning-Novel LMM

To more directly test the hypothesis that sentence processing after overlearning involves more activity in sensorimotor regions and less activity in ‘language regions’ compared to typical sentence processing, we did the same LMM as in the prior section but after first subtracting the coefficients for novel from overlearned sentences. We present the effects of each session separately because of the difficulty of interpreting the sign for this more complicated comparison. GLTs for overlearned minus novel sentences in both sessions, independent of time, show that overlearned sentences resulted in greater activity in sensorimotor regions. In contrast, novel sentences produce more activity in the superior temporal plane and inferior frontal gyrus in both sessions (Figure S3A and S3B). Subcortically, in session one, overlearned sentences produced more brainstem, nucleus accumbens and dorsal cerebellar activity while novel sentences produced more hippocampal activity. In session two, overlearned sentences produced more thalamus and more dorsal cerebellum activity while novel sentences produced more ventral cerebellar activity.

There were interactions between session and time for overlearned minus novel sentences that involved much of the brain though we do not attempt to interpret these here (though Figures 3 and 5 together suggest the direction of these effects). Nonetheless, to visualise changes over time as is Figure 3, the GLTs for each time point are presented separately for overlearned minus novel sentences in session one (Figure 5A) and session two (Figure 5B). The pattern of results clearly shows more sensorimotor activity for overlearned sentences in both sessions (Figure 5A and 5B, reds). The speech production LMM shows that this occurs in similar sensorimotor regions used to produce novel sentences. Conversely, there was less superior temporal plane and inferior frontal gyrus activity for overlearned sentences (Figure 5A and 5B, blues). These regions closely overlapped those for novel sentences alone (Figure 5, dark grey).

### Regional Homogeneity

The deconvolution and subsequent LMM results are model based, aggregating over stimuli, though without a priori assumptions about the shape of the hemodynamic response. To test whether there is support for hypotheses in a more model-free manner and to test whether there is more local sensorimotor connectivity after overlearning, we conducted a regional homogeneity analysis on preprocessed timeseries. There was an increase in sensorimotor cortices and a decrease in superior temporal plane local connectivity from session one to two for overlearned sentences (among other regions; Figure 6, top). We then directly contrasted overlearned and novel sentences for sessions one and two separately (Figure 6, bottom). In session one, there were few regions more active for overlearned sentences whereas novel sentences resulted in significantly greater regional connectivity, mostly throughout superior temporal plane and inferior parietal regions, bilaterally. In contrast, the overlearned sentences produced greater mostly inferior parietal and bilateral sensorimotor local connectivity in session two and the local superior temporal plane connectivity remained greater for novel sentences (Figure 6, bottom).

**Figure 6.**
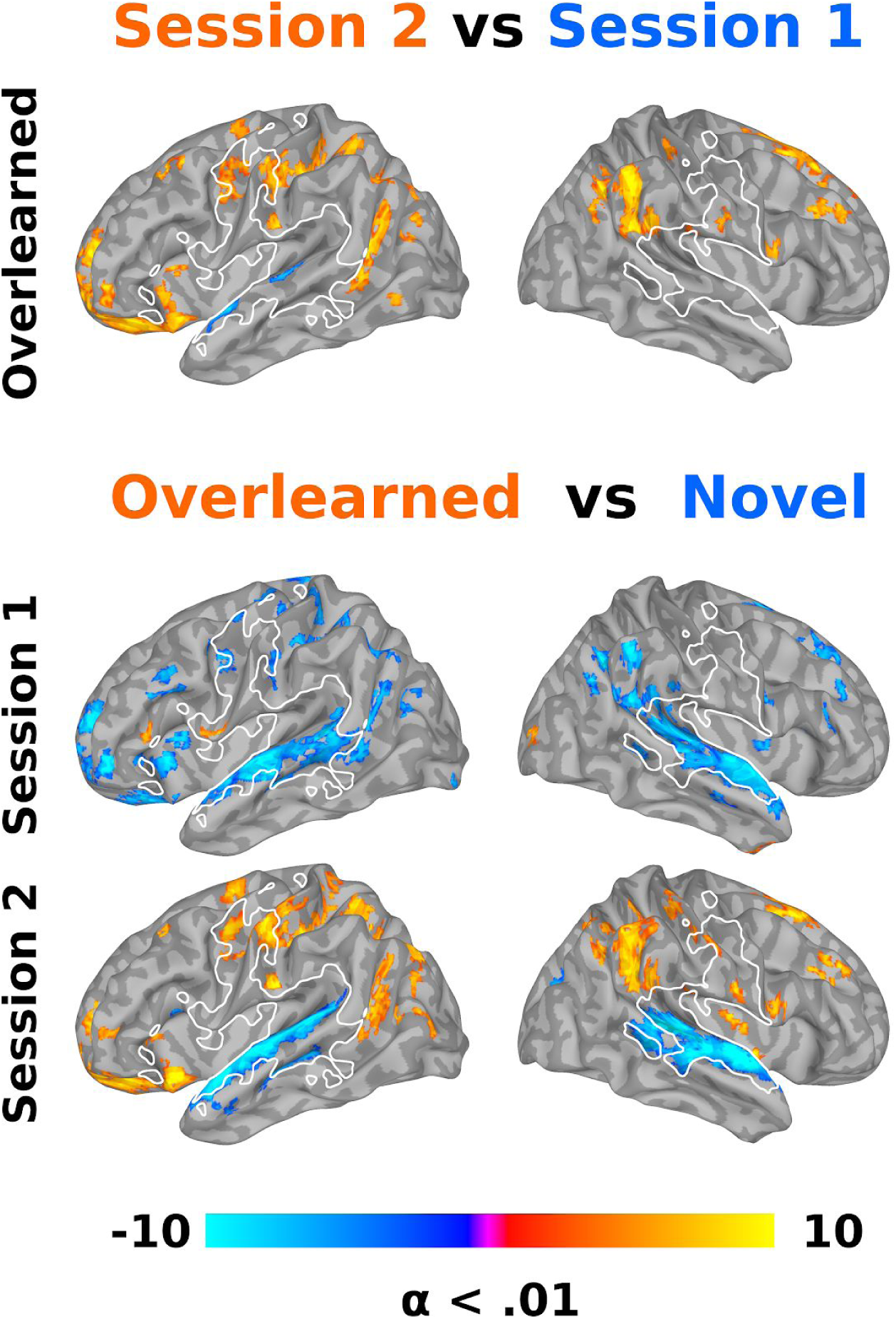
Local connectivity changes after overlearning. A regional homogeneity analysis was conducted to provide a more data driven validation of more model based results (shown in Figures 3-5) and to estimate changes in local synchronization or connectivity. In particular, we directly contrasted overlearned sentences between sessions one (blues) and two (reds; top row). We also contrasted novel (blues) and overlearned sentences (reds) for sessions one (middle row) and two (bottom row). Results confirm that listening to overlearned sentences after learning results in significantly more local connectivity in sensorimotor regions (reds) and significantly less local connectivity in the superior temporal plane.

### Network

To further test the hypothesis that the brain reorganises after overlearning, we analysed the global network variation between session one and two for overlearned sentences using edit distance. The average change in connectivity was 45.4% (SD = 5.50%; Minimum = 32.26%; Maximum = 55.37%), significantly different from the null hypothesis that there was no change from session one to two (t(11) = 28.69, p = 1.08 x 10^−11^). To visualise some of these changes in connectivity, we did chi-square (χ^2^) tests on the biniarised connections, using a threshold of 6.63 (p < .01; Figure 7). This resulted in 90 changes in connections, with 25 connections gained (27.77%) and 65 connections lost (72.22%). This large reduction in connections was across all major brain subdivisions. Nonetheless, a full 80% of the changes included medial / midline (42) and/or subcortical structures (34, with four overlapping medial-subcortical connections). Of the 42 medial regions, 27 were lost connections (64.29%). Of the 34 subcortical regions, 28 were lost connections (82.35%). New subcortical connections involved the amygdala (1x), cerebellum (1x), pallidum (1x), hippocampus (2x) and nucleus accumbens (2x). Lost subcortical connectivity involved the caudate (1x), amygdala (2x), cerebellum (2x), hippocampus (2x), pallidum (2x), brainstem (4x), putamen (5x), ventral diencephalon (6x) and thalamus (6x). At a more stringent threshold (p < .005), the disproportionate number of lost connections remains similar at 75.93%, with a disproportionate number of lost connections containing medial and subcortical connections (85.19%; see Table S1 for more specific information about changes in connectivity).

**Figure 7.**
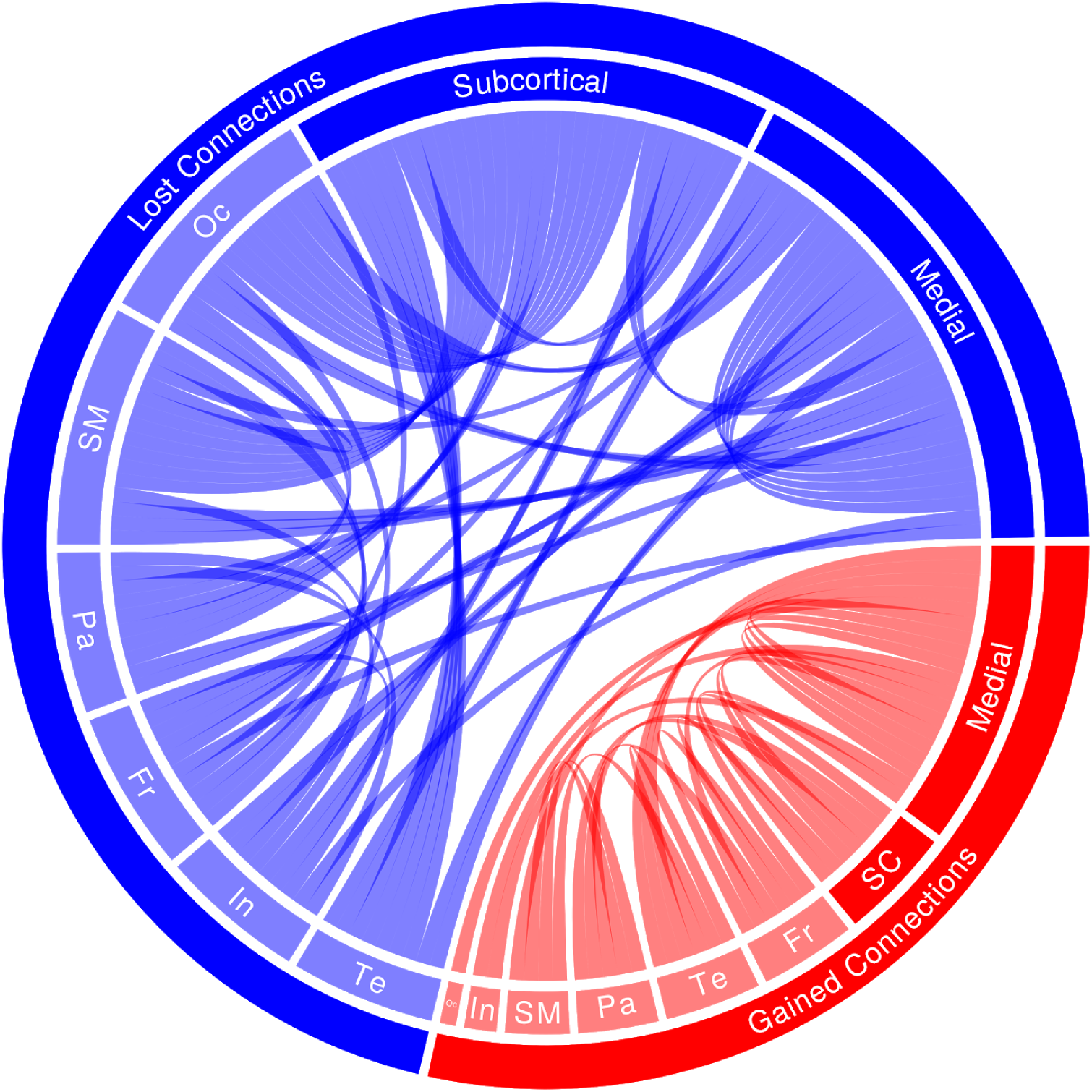
Changes in network connectivity after overlearning. The chord plot shows all 90 lost (blue) and gained (red) connections between sessions one and two as determined by chi-square tests on binarised connections between pairs of 167 regions of interest (χ^2^ > 6.63; p < .01). For ease of visualisation, these were grouped into frontal (Fr), insula (In), medial, occipital (Oc), parietal (Pa), sensorimotor (SM), subcortical (SC) and temporal (Te) regions. Results suggest that sentence overlearning results in significantly less connectivity, particularly in medial cortical and subcortical regions.

To characterise global connectivity changes with a more robust method, Deltacon similarity was computed. The average dissimilarity was 46.1%, (SD = 3.90%; Minimum = 43.20%; Maximum = 52.33%), with a significant change between sessions one and two (t(11) = 41.43, p = 1.97 x 10^−13^). To try and further understand network differences between sessions, we explored a number of other global and local measures. None of density, efficiency, diffusion or flow were significantly different from sessions one and two (ps > .05). There were also no differences between measures of centrality or community structure (ps > .05).

## Discussion

### Summary

We had participants overlearn sentences (Figure 1; Table 1) in an effort to evaluate the hypothesis that, as speech becomes more formulaic, it is processed more by sensorimotor cortices and not ‘language regions’. Behavioural results suggest that participants overlearned sentences. Reaction times to complete the final word of overlearned sentences decreased by 2.1 seconds (Figure 2A), participants’ identification of words drawn from overlearned sentences was faster (Figure 2B) and overlearned sentences were remembered with high accuracy (Figure 2C).

Correspondingly, there was a considerable and local increase after overlearning in sensorimotor regions centred around the central sulcus, including primary motor and somatosensory cortices in both model based and model-free analyses (Figures 3, 5 and 6, oranges). These regions overlapped with those involved in producing novel sentences (Figures 3, 5 and 6, white outline). After overlearning, speech perception begins earlier in sensorimotor regions, before the hemodynamic response is typically expected to rise (Figures 3 and 5, left) and before superior temporal regions (compare Figure 4, left and middle onsets). Consistent with reorganisation, some of these motor regions had a hemodynamic response that was centred around zero before overlearning (Figure 4, left).

Concomitant with these increases, we found a dramatic bilateral reduction in activity and local processing in the superior temporal plane when comparing overlearned sentences from sessions one to two (Figures 3, blues; Figure 6, blues, top). When overlearned sentences are compared to novel sentences, there was a reduction in activity and local processing in superior temporal plane and the inferior frontal gyrus ‘language regions’ (Figure 5, blues and grey; Figure 6, blues, bottom). Consistent with reorganisation, some superior temporal plane regions had a hemodynamic response around zero after overlearning, suggesting they were not active on average (Figure 4, middle).

Global network measures indicate that about 45% of the brain reorganised after overlearning. Much of this reorganisation involves a small number of new connections and a profound drop in the number of connections across the whole brain (Figure 7). The latter most saliently involved medial cortical and subcortical brain regions. Lost subcortical connections involved the amygdala, basal ganglia (mostly the putamen but also the caudate), brainstem, cerebellum, hippocampus, thalamus and ventral diencephalon. These subcortical results parallel those in voxel-based analyses that showed less increases and more decreases in activity (e.g., Figure 4, right).

### Overview

Results suggest that the brain mechanisms associated with perceiving overlearned speech are qualitatively different from less formulaic or more compositional language. They suggest that when speech segments become sufficiently overlearned, they are processed by a much more circumscribed and cortically isolated set of sensorimotor regions involved in producing speech and little, if at all, by ‘language regions’. As reviewed in the Introduction, the hypothesis predicting these results was predicated on 1) the relative preservation of both sensorimotor regions and formulaic language in aphasia and Alzheimer’s disease^45–49,63,65,66,75^ and 2) predictive models of the role of speech production related regions in speech perception^12,79,101^. Both suggest an increased role of sensorimotor and a decreased role of ‘language regions’ as learning increases. We review studies of formulaic language and motor sequence learning that mostly supports this proposal. Finally, we discuss implications for neurobiological models of language and therapy.

## Formulaic Language

### Production

It has been proposed that formulaic language production is supported by right hemisphere and subcortical interactions^45^. This conclusion was based on research showing that formulaic expressions are more common in left compared to right hemispheric damage^51–54^ (though see^49,133^). It was additionally based on results suggesting that individuals with Alzheimer’s disease produce more formulaic language than people with basal ganglia strokes^54,134^ and Parkinson’s disease^66,74^. This is attributed to relative preservation of the basal ganglia in Alzheimer’s disease. A region of interest based neuroimaging study in healthy people supports both arguments, showing that increased formulaic language production is correlated with increased right inferior frontal gyrus and decreased left caudate activity^135^.

Though we show some similar right hemisphere and subcortical effects, our results are more bilateral and centred more around sensorimotor rather than subcortical regions. To the extent our results can generalise to formulaic language production, they suggest that the postulated right hemisphere locus is due to a stronger weighting of right hemisphere regions after aphasia. Our results also suggest that the presumed subcortical locus might be less about the preservation or deterioration of the basal ganglia in Alzheimer’s and Parkinson’s disease and more about preserved sensorimotor regions in the former. This is more consistent with research suggesting that the basal ganglia is significantly impacted in Alzheimer’s disease, even in early stages^136–138^. Collectively, our results suggest that the more preserved right sensorimotor and not necessarily right ‘language regions’ or subcortical structures are the locus of formulaic speech production in aphasia and Alzheimer’s disease.

## Comprehension

Our results are more consistent with neuroimaging studies of formulaic related language comprehension. Much of this work centres around figurative language, like idioms and metaphors. These are processed bilaterally in ‘language regions’, with varying contribution of the left and right hemisphere as a function of, among other things, familiarity^139–145^. Furthermore, the more familiar^146–152^, frequent^153–157^, compositional (‘red boat’ vs ‘cup boat’) or coherent multiword expressions are^158,159^, the less ‘language regions’ tend to be active. Some studies show that sensorimotor activity is more strongly associated with high frequency words, word composition and impairments of word composition^160,161^ while the basal ganglia are less active for higher frequency words^157,162^. One study concluded that ‘areas canonically implicated in traditional neurophysiological models of language processing appear to play a lesser role in basic composition’ (p. 2802)^160^. Another concluded that multiword expressions rely on regions other than ‘traditional frontal and temporal nodes of the language network’ (p. 12)^158^.

### Sequence Learning

The strongest similarities to our results derive from motor sequence learning research. This work shows that becoming or being an expert motor performer involves a well documented set of decreases and increases in brain activity that depend on the length of learning^163–173^. Specifically, there are two learning stages associated with relative duration: fast online learning, described as more explicit and by repetition suppression and slow learning, described as more implicit and by repetition enhancement and sleep related or offline consolidation. Fast learning is frequently linked to more associative cortical and subcortical regions. In contrast, slow learning is associated with a global decrease in activity in most regions, including prefrontal, premotor, parietal, sensorimotor and subcortical structures like more associative basal ganglia and cerebellar regions. Amongst these widespread decreases, there is a selective increase in some sensorimotor and subcortical regions, including less associative aspects of the basal ganglia and cerebellum. Generally, these changes might be described as a shift away from cognitive systems (involving attentional, inhibition, control, etc.) and towards more ‘automatic’ sensorimotor brain regions.

Our study incorporated early and late learning phases, with an early online perceptual learning period (measured in fMRI session one) and a late offline period of production based learning with motor memory consolidation (measured in fMRI session two). Consistent with slow learning, we show global decreases in activity in most cortical and subcortical regions with a selective focal increase in some sensorimotor and subcortical regions. Also consistent with the distinction between fast and slow learning, our results do not simply constitute a redistribution of activity patterns in the same ‘language regions’ but, rather, are a reorganization to sensorimotor regions, suggesting different processes underlying the perception of novel and overlearned sentences^165^.

### Neurobiological Models

More generally, our results suggest a model by which the perception of overlearned speech may not rely on ‘language regions’ or ‘the language network’. Like test-retest reliability and lexical processing studies (discussed in the Introduction), our results suggest that the neurobiology of speech perception and language comprehension is more variable and distributed than posited by classical and contemporary models^1–11^.

If so, an open question becomes what differentiates the more distributed regions found in some studies from the fixed set of regions in popular models? One hypothesis is that the whole brain variously participates in speech perception and language comprehension and that ‘language regions’ as we know them are mostly connectivity hubs coordinating this more distributed system^12^. Because of the reliance on measures of central tendency, these distributed regions are averaged out in most studies^12^. They only become obvious when specific categories are under examination, e.g., individual differences, action words or, as here, formulaic language.

Indeed, most ‘language regions’ are structural and functional connectivity hubs in humans^174–177^, as are homologous regions in monkeys^178–181^. As befitting connectivity hubs, ‘language regions’ are some of the most ‘reused’ regions in the human brain, participating in many perceptual, motor and cognitive domains beyond language^182^. Given how central and diverse they are, it is perhaps not surprising that damage to these hubs predict neuropsychological outcomes beyond language problems^183,184^.

If they are mostly hubs, the focus on ‘language regions’ or their ‘homologues’ in therapeutic interventions runs the risk of overemphasizing the importance of less specific regions and neglecting more behaviourally relevant network nodes. For example, given the preservation of formulaic language and corresponding sensorimotor regions in some individuals with aphasia and Alzheimer’s disease, it makes sense to focus on those expressions and regions as they might be used as a scaffold for language recovery. Indeed, use of formulaic song and language shows promise in therapy^185–190^. In contrast, the relative preservation of some other aspect of language and associated distributed set of network nodes might benefit from a different intervention.

To the extent models guide therapy, this proposal requires us to move beyond current neurobiological models with static regions^2,12,79,191,192^. It begs for a more detailed neurobiological account of overlearned expressions and other factors that result in differently distributed language processes. It also suggests a greater focus on item analysis and individual differences and a less strong reliance on measures of central tendency to understand the organisation of language and the brain^193^.

## Limitations

Overlearning often occurs through repetition in short sessions over a relatively small period of time (e.g., as in learning at home over a period of weeks for school tests). However, much of formulaic language is picked up over a lifetime of repetitions. Similarly, though many overlearned expressions are not particularly meaningful (e.g., ‘whats up’), most do have more specific meanings. Our repetition learning task did not unfold over years nor did it emphasise semantic content per se. Thus, the neurobiology of more extensive overlearning, with more semantically meaningful content might differ from what was observed here.

As with any longitudinal design, it might be argued that participants were not attending to overlearned stimuli or attending as much in session two. Participants were monitored for wakefulness during scanning and all appeared to be attending in both sessions. Post scan recall indicates that participants paid attention enough to recall novel sentences with relatively high accuracy.

## Conclusions

Results suggest that the brain regions supporting speech perception are not fixed but, rather, dramatically reorganise as a function of individual experience with speech production. Specifically, repeated experience speaking the same word sequence seems to change the memory representation of those words to be more formulaic. This trace is subsequently used by the brain in the process of speech perception, perhaps in a predictive manner. This involves a different set of regions than is said to support more compositional language. Given how frequently formulaic expressions occur, how fast they are processed and important they are in learning, results call for more research on how overlearning occurs and formulaic speech is processed. They also suggest why formulaic expressions might be preserved in aphasia and Alzheimer’s disease. Therapy for such disorders, if they are neurobiologically informed, are presumably informed by prevailing beliefs. One of these is the (often implicit) belief that language is supported by a static set of ‘language regions’ or ‘the language network’, formalised in classical and contemporary neurobiological models. Given that this is not the case, as shown here and elsewhere, future interventions might benefit from adopting more dynamic and distributed network models of language and the brain^12^.

## Acknowledgments

JIS would like to thank the fine people at the Birkbeck-UCL Centre for Neuroimaging (BUCNI) and acknowledge support from EPSRC EP/M026965/1 ‘New pathways to hearing: A multisensory noise reducing and palate based sensory substitution device for speech perception.’ JIS would like to thank Marple and BB. Hair are your aerials. DRL would like to acknowledge support from the British Academy and the Natural Sciences and Engineering Research Council of Canada.

## Author Contributions

DL and JIS conceived of the study. SB, DL, EM and JIS designed the study. All authors helped with stimulus pretesting and EM selected overlearned sentences. JIS wrote the manuscript with help from SA, SB, YJJ and DL. EM, SB, SL and YJJ collected the data. DL did behavioural analyses. SL and YJJ did the ICA artifact labelling for fMRI preprocessing, checked by JIS. JIS did all fMRI preprocessing and analyses except the network analyses, done by SA.

## Data Availability

All fMRI data will be made available on OpenNeuro (https://openneuro.org/). Scripts used to analyse data will be made available on GitHub (https://github.com/lab-lab). Images used to make all figures will be made available on NeuroVault (https://neurovault.org/).

