## Supplementary Material for "Reorganization of the neurobiology of language after sentence overlearning"

#### Figure Captions

**Figure S1.** Additional novel listening linear mixed-effects model results. General linear test contrasting novel sentences from session one (blues) and two (reds) in the left (LH) and right hemispheres (RH) presented on lateral (top) and medial (bottom) surface views. The colour bar represents z-scores and the images are thresholded at an alpha (α) level of $p < .01$, corrected for multiple comparisons.

**Figure S2.** Additional overlearned sentence listening linear mixed-effects model results. A) Session one general linear test (GLT) for sentence; B) Session two GLT for sentence; C) Direct contrast of session one sentences (blues) with session two sentences (reds). The top two rows are the left hemisphere (LH) while the bottom two are the right hemispheres (RH) lateral and medial surface views. The colour bar represents z-scores and the images are thresholded at an alpha (α) level of $p < .01$, corrected for multiple comparisons.

**Figure S3.** Additional overlearned minus novel sentence listening linear mixed-effects model results. A) Session one general linear test (GLT) for overlearning-novels sentences and B) Session two GLT for overlearning-novels sentences. Colours indicate novel sentences (blues) and overlearned sentences (reds). The top two rows are the left hemisphere (LH) while the bottom two are the right hemispheres (RH) lateral and medial surface views. The colour bar represents z-scores and the images are thresholded at an alpha (α) level of $p < .01$, corrected for multiple comparisons.

*Figure S1*

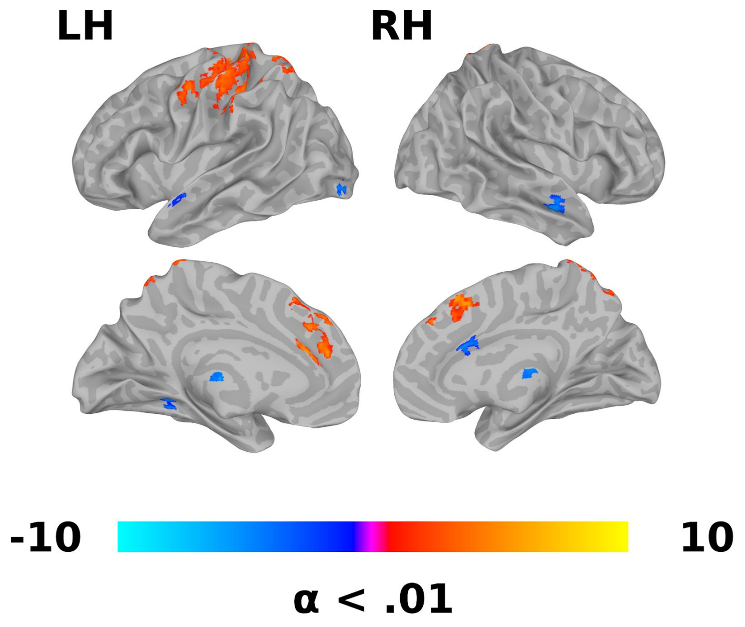

*Figure S2*

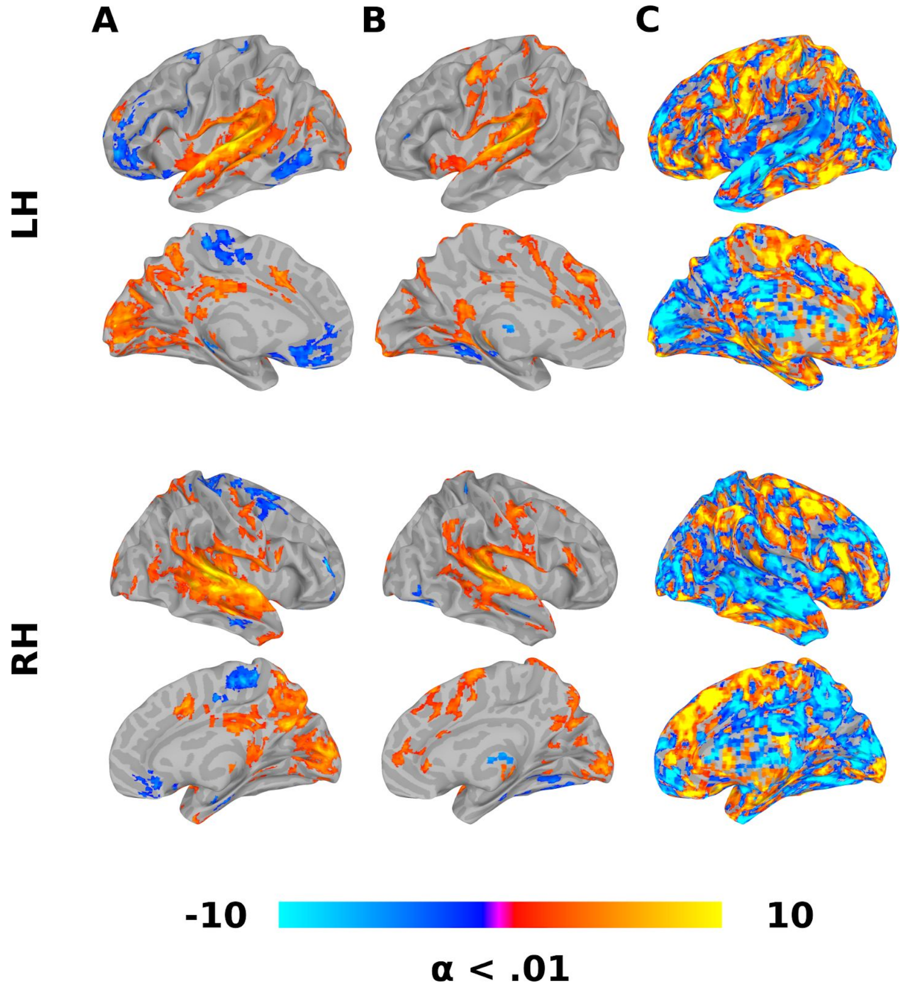

***Figure S3***

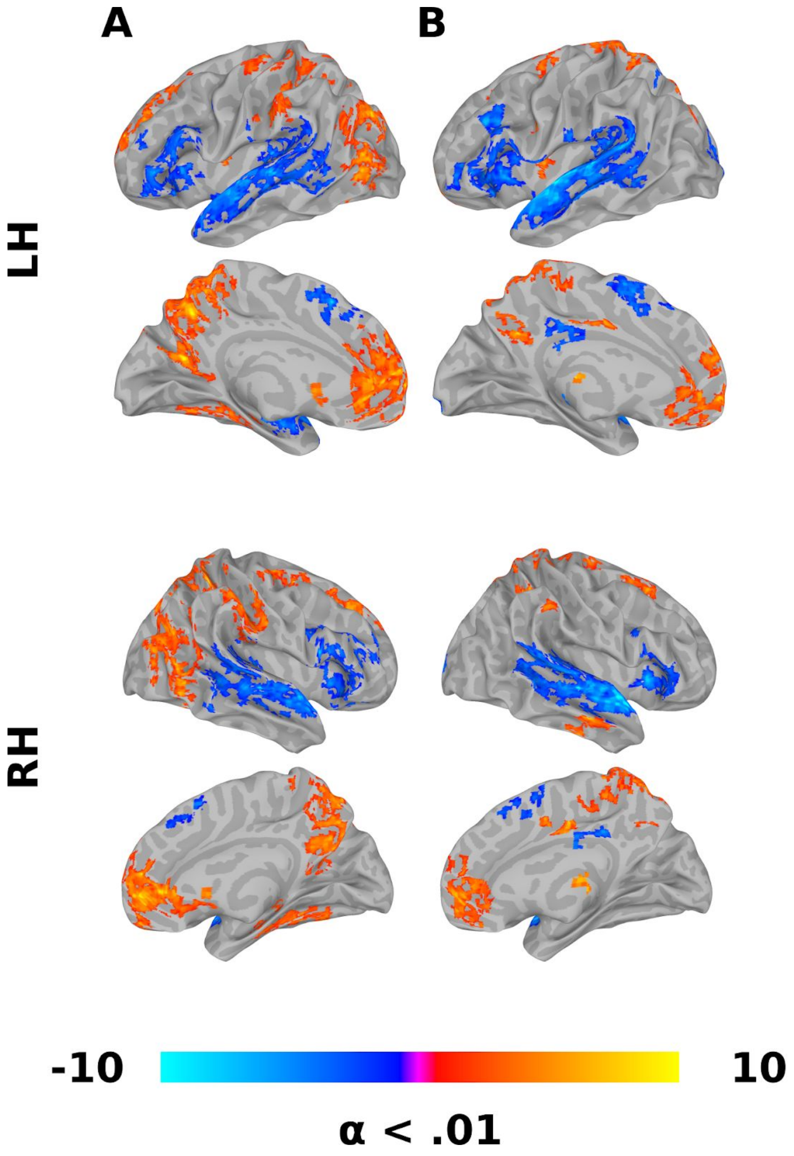

### Tables

**Table S1.** Change in network connections from session one and two.

| | Gross Regions | | Specific Regions | | |
|---|---|---|---|---|---|
| Change | Source | Target | Source | Target | P-Value |
| Loss | Frontal | Medial | LH Inferior Frontal Sulcus | RH Transverse Frontopolar Gyri and Sulci | 0.009 |
| Loss | Frontal | Occipital | RH Middle Frontal Gyrus | RH Occipital Pole | 0.005 |
| Loss | Frontal | Parietal | RH Superior Frontal Sulcus | RH Intraparietal Sulcus and Transverse Parietal Sulci | 0.009 |
| Loss | Frontal | Sensori motor | LH Orbital Sulci | RH Precentral Gyrus | 0.003 |
| Loss | Frontal | Sensori motor | RH Vertical Ramus of the Anterior Lateral Fissure | RH Central Sulcus | 0.005 |
| Loss | Insula | Frontal | LH Anterior Segment of the Circular Sulcus of the Insula | LH Orbital Sulci | 0.009 |
| Loss | Insula | Medial | RH Inferior Segment of the Circular Sulcus of the Insula | RH Subparietal Sulcus | 0.000 |
| Loss | Insula | Parietal | RH Long Insular Gyrus and Central Sulcus of the Insula | RH Precuneus | 0.009 |
| Loss | Medial | Frontal | LH Transverse Frontopolar Gyri and Sulci | LH Inferior Frontal Sulcus | 0.001 |
| Loss | Medial | Frontal | RH Subcallosal Gyrus | RH Orbital Sulci | 0.003 |
| Loss | Medial | Insula | LH Parahippocampal Gyrus | LH Inferior Segment of the Circular Sulcus of the Insula | 0.005 |
| Loss | Medial | Insula | LH Anterior Transverse Collateral Sulcus | RH Anterior Segment of the Circular Sulcus of the Insula | 0.009 |
| Loss | Medial | Medial | LH Transverse Frontopolar Gyri and Sulci | LH Lateral Orbital Sulcus | 0.005 |
| Loss | Medial | Medial | LH Orbital Gyrus | RH Subcallosal Gyrus | 0.005 |

| | | | | | |
|---|---|---|---|---|---|
| Loss | Medial | Medial | LH Marginal Branch of the Cingulate Sulcus | RH Suborbital Sulcus | 0.009 |
| Loss | Medial | Medial | LH Medial Orbital Sulcus | RH Subparietal Sulcus | 0.009 |
| Loss | Medial | Medial | RH Marginal Branch of the Cingulate Sulcus | RH Suborbital Sulcus | 0.009 |
| Loss | Medial | Occipital | LH Frontomarginal Gyrus and Sulcus | LH Fusiform Gyrus | 0.001 |
| Loss | Medial | Occipital | LH Superior Frontal Gyrus | LH Middle Occipital Gyrus | 0.004 |
| Loss | Medial | Parietal | RH Posterior Ventral Cingulate Gyrus | RH Angular Gyrus | 0.003 |
| Loss | Medial | Sensori motor | RH Subcallosal Gyrus | RH Central Sulcus | 0.009 |
| Loss | Medial | Tempor al | LH Subparietal Sulcus | RH Transverse Temporal Gyrus | 0.004 |
| Loss | Medial | Tempor al | LH Subparietal Sulcus | RH Transverse Temporal Sulcus | 0.004 |
| Loss | Occipital | Insula | LH Occipital Pole | RH Long Insular Gyrus and Central Sulcus of the Insula | 0.009 |
| Loss | Occipital | Medial | LH Occipital Pole | RH Anterior Transverse Collateral Sulcus | 0.002 |
| Loss | Occipital | Medial | LH Middle Occipital Gyrus | RH Posterior Ventral Cingulate Gyrus | 0.003 |
| Loss | Occipital | Parietal | LH Middle Occipital Gyrus | RH Superior Parietal Lobule | 0.001 |
| Loss | Occipital | Parietal | LH Anterior Occipital Sulcus and Preoccipital Notch | RH Superior Parietal Lobule | 0.004 |
| Loss | Occipital | Sensori motor | LH Medial Occipito-Temporal Sulcus and Lingual Sulcus | RH Paracentral Gyrus and Sulcus | 0.001 |
| Loss | Parietal | Frontal | RH Supramarginal Gyrus | RH Inferior Frontal Sulcus | 0.009 |
| Loss | Parietal | Medial | LH Superior Parietal Lobule | RH Marginal Branch of the Cingulate Sulcus | 0.009 |

| | | | | | |
|---|---|---|---|---|---|
| Loss | Sensori motor | Insula | LH Paracentral Gyrus and Sulcus | RH Short Insular Gyri | 0.004 |
| Loss | Sensori motor | Medial | RH Subcentral Gyrus and Sulcus | RH Anterior Transverse Collateral Sulcus | 0.005 |
| Loss | Sensori motor | Medial | LH Postcentral Gyrus | LH Parieto-Occipital Sulcus | 0.009 |
| Loss | Sensori motor | Medial | RH Subcentral Gyrus and Sulcus | RH Pericallosal Sulcus | 0.009 |
| Loss | Subcorti cal | Frontal | LH Pallidum | LH Vertical Ramus of the Anterior Lateral Fissure | 0.009 |
| Loss | Subcorti cal | Insula | LH Cerebellum | RH Long Insular Gyrus and Central Sulcus of the Insula | 0.001 |
| Loss | Subcorti cal | Insula | LH Ventral Diencephalon | LH Inferior Segment of the Circular Sulcus of the Insula | 0.001 |
| Loss | Subcorti cal | Medial | LH Thalamus | LH Superior Frontal Gyrus | 0.004 |
| Loss | Subcorti cal | Medial | RH Thalamus | LH Anterior Cingulate Gyrus and Sulcus | 0.009 |
| Loss | Subcorti cal | Occipital | RH Putamen | RH Anterior Occipital Sulcus and Preoccipital Notch | 0.001 |
| Loss | Subcorti cal | Occipital | LH Cerebellum | RH Calcarine Sulcus | 0.003 |
| Loss | Subcorti cal | Occipital | Brainstem | LH Anterior Occipital Sulcus and Preoccipital Notch | 0.003 |
| Loss | Subcorti cal | Occipital | Brainstem | RH Cuneus | 0.009 |
| Loss | Subcorti cal | Occipital | Brainstem | RH Superior Occipital Gyrus | 0.009 |
| Loss | Subcorti cal | Occipital | RH Pallidum | RH Calcarine Sulcus | 0.009 |
| Loss | Subcorti cal | Parietal | RH Putamen | RH Superior Parietal Lobule | 0.001 |

| | | | | | |
|---|---|---|---|---|---|
| Loss | Subcortical | Parietal | LH Thalamus | RH Sulcus Intermedius Primus of Jensen | 0.004 |
| Loss | Subcortical | Parietal | RH Ventral Diencephalon | RH Sulcus Intermedius Primus of Jensen | 0.009 |
| Loss | Subcortical | Sensorimotor | LH Amygdala | RH Precentral Gyrus | 0.003 |
| Loss | Subcortical | Sensorimotor | LH Thalamus | RH Superior Precentral Sulcus | 0.004 |
| Loss | Subcortical | Sensorimotor | LH Amygdala | LH Paracentral Gyrus and Sulcus | 0.004 |
| Loss | Subcortical | Sensorimotor | RH Caudate | RH Superior Precentral Sulcus | 0.004 |
| Loss | Subcortical | Sensorimotor | RH Hippocampus | RH Paracentral Gyrus and Sulcus | 0.004 |
| Loss | Subcortical | Sensorimotor | LH Thalamus | LH Postcentral Sulcus | 0.009 |
| Loss | Subcortical | Sensorimotor | LH Ventral Diencephalon | RH Postcentral Gyrus | 0.009 |
| Loss | Subcortical | Subcortical | LH Thalamus | RH Hippocampus | 0.009 |
| Loss | Subcortical | Subcortical | Brainstem | RH Putamen | 0.009 |
| Loss | Subcortical | Temporal | LH Ventral Diencephalon | LH Transverse Temporal Gyrus | 0.004 |
| Loss | Subcortical | Temporal | LH Ventral Diencephalon | LH Posterior Lateral Fissure | 0.004 |
| Loss | Subcortical | Temporal | LH Ventral Diencephalon | RH Transverse Temporal Sulcus | 0.004 |
| Loss | Subcortical | Temporal | RH Putamen | LH Inferior Temporal Sulcus | 0.004 |
| Loss | Subcortical | Temporal | RH Putamen | RH Inferior Temporal Gyrus | 0.004 |

| | | | | | |
|---|---|---|---|---|---|
| Loss | Temporal | Medial | RH Superior Temporal Gyrus | RH Pericallosal Sulcus | 0.002 |
| Loss | Temporal | Medial | LH Inferior Temporal Gyrus | RH Pericallosal Sulcus | 0.005 |
| Gain | Frontal | Frontal | RH Horizontal Ramus of the Anterior Lateral Fissure | RH Superior Frontal Sulcus | 0.009 |
| Gain | Frontal | Medial | LH Pars Orbitalis | LH Marginal Branch of the Cingulate Sulcus | 0.009 |
| Gain | Insula | Medial | LH Anterior Segment of the Circular Sulcus of the Insula | RH Lateral Orbital Sulcus | 0.002 |
| Gain | Medial | Frontal | LH Suborbital Sulcus | RH Middle Frontal Sulcus | 0.004 |
| Gain | Medial | Insula | LH Transverse Frontopolar Gyri and Sulci | RH Long Insular Gyrus and Central Sulcus of the Insula | 0.009 |
| Gain | Medial | Medial | LH Superior Frontal Gyrus | LH Subcallosal Gyrus | 0.001 |
| Gain | Medial | Medial | LH Middle-Anterior Cingulate Gyrus and Sulcus | LH Subcallosal Gyrus | 0.003 |
| Gain | Medial | Medial | LH Suborbital Sulcus | RH Middle-Anterior Cingulate Gyrus and Sulcus | 0.003 |
| Gain | Medial | Medial | RH Middle-Anterior Cingulate Gyrus and Sulcus | RH Pericallosal Sulcus | 0.005 |
| Gain | Medial | Occipital | LH Anterior Cingulate Gyrus and Sulcus | LH Lingual Gyrus | 0.003 |
| Gain | Medial | Sensorimotor | LH Subcallosal Gyrus | LH Superior Precentral Sulcus | 0.004 |
| Gain | Medial | Temporal | LH Superior Frontal Gyrus | RH Middle Temporal Gyrus | 0.004 |
| Gain | Medial | Temporal | LH Lateral Orbital Sulcus | RH Inferior Temporal Gyrus | 0.005 |
| Gain | Medial | Temporal | LH Lateral Orbital Sulcus | RH Superior Temporal Sulcus | 0.009 |
| Gain | Parietal | Frontal | RH Precuneus | RH Vertical Ramus of the Anterior Lateral Fissure | 0.009 |

| | | | | | |
|---|---|---|---|---|---|
| Gain | Parietal | Sensorimotor | LH Sulcus Intermedius Primus of Jensen | RH Postcentral Gyrus | 0.004 |
| Gain | Parietal | Sensorimotor | LH Sulcus Intermedius Primus of Jensen | RH Central Sulcus | 0.009 |
| Gain | Subcortical | Frontal | LH Nucleus Accumbens | RH Pars Triangularis | 0.009 |
| Gain | Subcortical | Medial | RH Hippocampus | LH Subcallosal Gyrus | 0.003 |
| Gain | Subcortical | Medial | RH Nucleus Accumbens | LH Middle-Posterior Cingulate Gyrus and Sulcus | 0.009 |
| Gain | Subcortical | Parietal | LH Hippocampus | LH Sulcus Intermedius Primus of Jensen | 0.009 |
| Gain | Subcortical | Parietal | LH Amygdala | RH Sulcus Intermedius Primus of Jensen | 0.009 |
| Gain | Subcortical | Subcortical | RH Cerebellum | RH Pallidum | 0.001 |
| Gain | Temporal | Sensorimotor | LH Planum Polare | LH Inferior Precentral Sulcus | 0.009 |
| Gain | Temporal | Temporal | LH Inferior Temporal Sulcus | RH Middle Temporal Gyrus | 0.009 |
